# Individualized and stereotypical seizure semiology in a porcine model of post-traumatic epilepsy

**DOI:** 10.64898/2026.02.26.708000

**Authors:** Marco Pretell, Mariana Gonzalez Rodriguez, Waizo Chen, Alejandro Escobosa, Nelson Marquez Carvajal, Luis Martinez Ramirez, Benjamin Baskin, Monica Tynan, Andrew Schwalb, Paige O’Gorman, Arihant Patel, Caitlyn Smith, Veronica Sendao, Justinne Quinanola, Rayen Gandhi, Aarush Patnala, Kyle P Lillis, Kevin Staley, Beth Costine-Bartell

**Affiliations:** Department of Neurosurgery, Massachusetts General Hospital, Charlestown, Massachusetts, United States of America; Department of Neurology, Massachusetts General Hospital, Charlestown, Massachusetts, United States of America; Department of Neurosurgery, Harvard Medical School, Boston, Massachusetts, United States of America; Department of Neurology, Harvard Medical School, Boston, Massachusetts, United States of America

**Keywords:** Post-traumatic epilepsy, semiology, cortical impact, swine, convulsions, seizure, early traumatic seizure

## Abstract

Large-animal models of traumatic brain injury may yield translatable data on epileptogenesis, given their similarities in anatomy, brain size, and immune systems to humans. Adult male and female swine received bilateral cortical impact (N=16) or sham surgery (N=6) and were screened for convulsions via video-EEG for up to one year. Post-traumatic epilepsy (PTE) was defined as 2 seizures after 1-week post-injury. Nine out of sixteen pigs (56%) receiving bilateral cortical impact developed PTE, with an average 4.6 months (±3.4,SD) latent period. Seizures (N = 199) began focally, sometimes with motor onset including automatisms, before becoming generalized, with tonic-clonic or tonic convulsions. We defined a library differentiating peri-ictal behaviors (N = 31) from rhythmic/odd behaviors typical in healthy pigs (N = 12). Seizures had an average of 7.3 behaviors per seizure (max 26) lasting an average of 1.8 minutes (max 7.9). For seizures comprised of multiple convulsive episodes, the first convulsion had a greater number of peri-ictal behaviors than subsequent convulsions (P < 0.001). The array of peri-ictal behaviors displayed was pig-specific, with many behaviors consistently observed across seizures. The seizure frequency detected was 0.38/day. This large-brain model of PTE exhibits a variable period of epileptogenesis, a substantial rate of PTE, and an expansive repertoire of ictal behaviors. This first description of semiology in this species will serve as a guide for other porcine epilepsy models. Biofidelic models of PTE are expected to increase our understanding of the pathophysiology, enabling the identification and testing of therapeutics that translate into human patients.

**Highlights:**

1. The average time from bilateral cortical impact to post-traumatic epilepsy in swine is 6 months, and is highly variable, ranging from 2 to 47 weeks post-TBI.
2. Swine with post-traumatic epilepsy display an array of specific behaviors around convulsions, distinct from pigs without post-traumatic epilepsy.
3. Though the duration of convulsion was typically a few seconds, the entire seizure, with the associated peri-ictal behaviors, lasts up to 7.9 minutes.
4. The complexity of behaviors around convulsions tended to increase from early traumatic seizures to post-traumatic seizures.
5. Peri-ictal behaviors observed around convulsions in an individual were often displayed prior to the first convulsion.

**Graphical Abstract:** 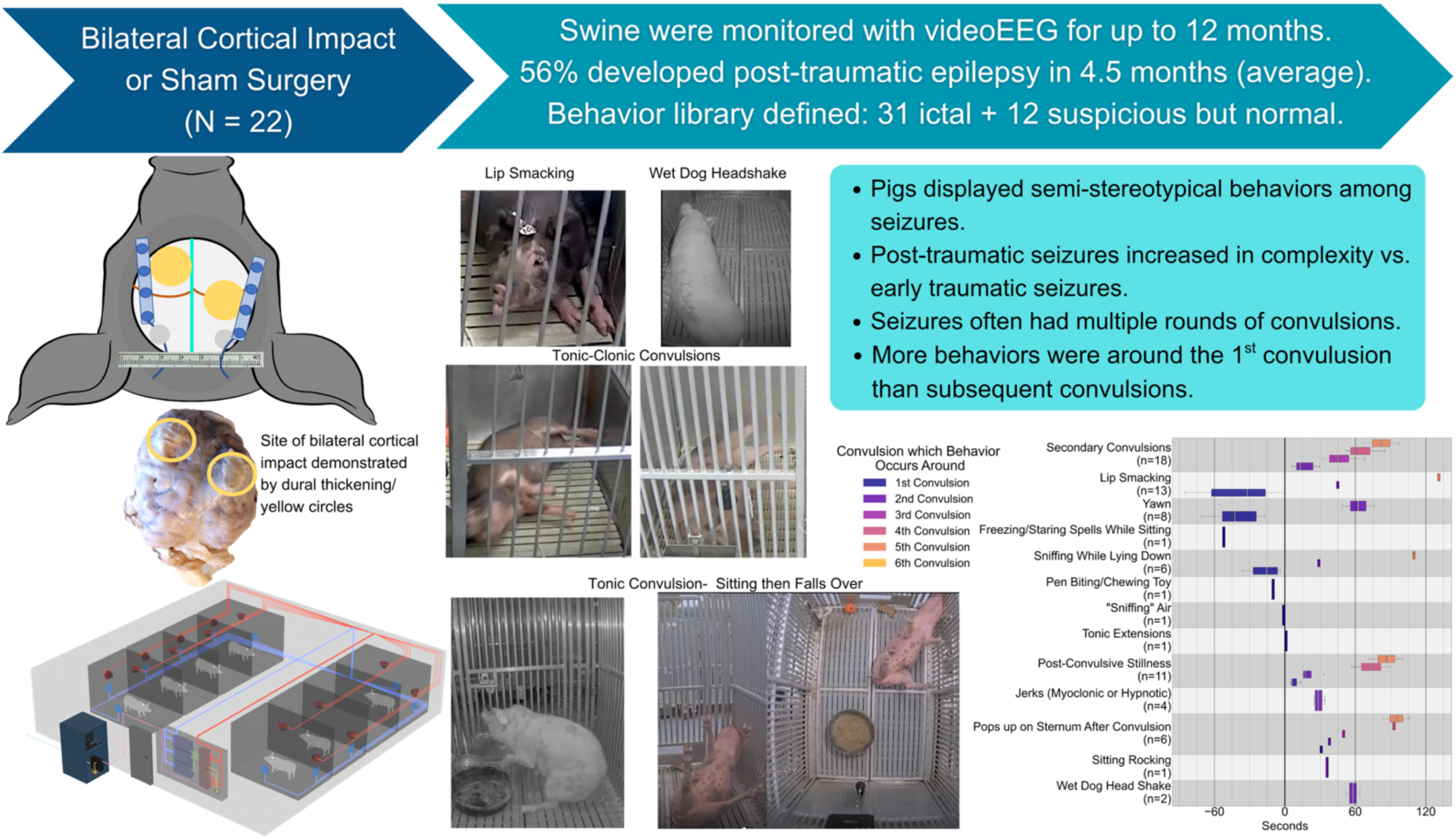

## Introduction

Post-traumatic epilepsy (PTE) can be a long-term consequence of traumatic brain injury (TBI), complicating or even slowing recovery (Verellen and Cavazos 2010), and is highly resistant to treatment with anti-epileptic drugs (Temkin, Dikmen et al. 1990, Garga and Lowenstein 2006). PTE is defined as spontaneous, recurrent seizures occurring at least one week after injury. The mechanisms underlying epileptogenesis remain unknown, in part, due to the prolonged latent period between injury and seizure onset, creating logistical challenges for studying the disease (Yazdi, Gomez et al. 2025).

Preclinical models are important for understanding the mechanisms underlying translationally relevant epileptogenesis. Rodent models of TBI-induced epilepsy have provided important insights, but are limited in their ability to replicate important characteristics of human PTE. Most rodent TBI models injure the cortex, which is thin, while also injuring the underlying hippocampus. This often results in pathology and seizure phenotypes that differ from epilepsy observed in human PTE, where most impacts and contusions are on the frontal or temporal lobes, not the hippocampus (Brady, Casillas-Espinosa et al. 2019). In addition, differences in brain size, cortical organization, white matter composition, glial responses, and immune signaling complicate translation from rodents to humans, particularly for disorders with long latent periods, such as PTE (Brady, Casillas-Espinosa et al. 2019). The latent period in mice (Reddy, Golub et al. 2025) is much shorter than the latent period in humans (Annegers, Grabow et al. 1980, Annegers, Hauser et al. 1998).

Swine share important anatomical features with humans, including cortical folding, a high proportion of white matter, and prolonged post-injury remodeling (Sauleau, Lapouble et al. 2009). Importantly, a larger brain size is associated with longer epileptogenic latency (Lillis, Wang et al. 2015), requiring long-term monitoring to capture seizure onset and progression. Recent advances in fully implantable, wireless electroencephalographic (EEG) systems have enabled long-term video-EEG monitoring in swine following bilateral cortical impact. While prior work has established the feasibility of inducing PTE and recording electrographic seizures in this model (Martinez-Ramirez, Slate et al. 2022), the rate of PTE and semiology in this model have not yet been characterized.

In both preclinical and clinical epilepsy research, seizure semiology is essential to determine seizure onset and propagation and epilepsy classification (Fisher, Cross et al. 2017, Beniczky, Tatum et al. 2022). Visual analysis of EEG alone is notoriously fraught with moderate inter-rater reliability among epileptologists (Halford, Shiau et al. 2015), and video-EEG is recommended (Tatum, Rubboli et al. 2018). Seizures in mice, even when induced via cortical impact and not a chemical convulsant, can have a brief latent period between early traumatic seizures and PTE (Williams, White et al. 2006, Shandra, Mahmutovic et al. 2026), and seizures are very brief, lacking the semiological complexity present in large-brained species, including humans (Racine 1972, Lüttjohann, Fabene et al. 2009, Williams, White et al. 2009). The present study addresses this translational gap by systematically characterizing seizure semiology in a porcine model of PTE following bilateral cortical impact using long-term, synchronized video electrocorticography (ECoG), positioning swine as a translational model that bridges rodent and human epilepsy and reflects both shared and species-specific features.

## Methods and Materials

### Animals and Housing

All procedures were approved by the MGH Institutional Care and Use Committee (protocol # 2018N000071: “Role of extracellular matrix injury in post-traumatic epilepsy”) in accordance with Federal Office of Lab Animal Welfare Guidelines and the American Veterinary Association Guidelines. All 10 elements of the Animal Research: Reporting of In Vivo Experiments Guidelines are addressed below. Castrated male and intact female adult Yucatan pigs were housed in a 12-hour light:12-hour dark cycle with lights on at 7:00 am and lights off at 7:00 pm. All pens were equipped with enrichment activities, and the animals received regular exercise sessions.

The animals described here were divided into two cohorts. In both cohorts, cortical impact was deployed as previously described through burr holes in the skull to the somatosensory cortex of the snout (rostral gyrus) on the right hemisphere and the very rostral portion of the left hemisphere (mean age = 6.9 ± 3.2 months (SD), mean weight = 36.4 ± 13.6 kg, SD)(Martinez-Ramirez, Slate et al. 2022). This array was intended to cause a pathologically significant lesion and a high rate of PTE, with the impacts offset to minimize clinical signs. Swine were generally somnolent the day after TBI, and some had an early traumatic seizure, but otherwise were not symptomatic until PTE developed. The anesthesia and surgical procedures have been previously described (Martinez-Ramirez, Slate et al. 2022). Briefly, pigs were anesthetized with Telazol, followed by isoflurane, and then switched to a mostly non-GABA acting sedation (dexmedetomidine, morphine, rocuronium). Non-steroidal anti-inflammatory medications were not used for analgesia so as not to interfere with pathophysiology, but analgesia was provided with local bupivacaine, buprenorphine (short- or long-acting), and/or fentanyl patches for at least 72 hours after each surgery and additionally as needed through breakthrough pain.

Cohort 1 was a pilot study involving a series of pigs (not randomized) that received either bilateral cortical impact (N = 10) or sham surgery (N = 3). A subset of these pigs (N = 3) also received AAV injection into the cortex for an additional experimental aim, performed a month before brain collection (Costine-Bartell, Martinez-Ramirez et al. 2023). In this cohort, EEG electrodes were attached to the skull via skull screws (Martinez-Ramirez, Slate et al. 2022), which was problematic as previously described (Martinez-Ramirez, Slate et al. 2022). None were excluded in Cohort 1.

Cohort 2 pigs were randomized to bilateral cortical impact (N = 6) or sham surgery (N = 3), and all had AAV injected into the cortex a month before cortical impact or sham surgery for a separate experimental question as previously described (Costine-Bartell, Martinez-Ramirez et al. 2023). Following the craniotomy, two EEG transceivers were placed subcutaneously in the neck and spliced to two silicone-platinum electrode strips (AdTech), which were placed epidurally to cover the dorsal cortex adjacent to the cortical impact or sham sites (**Figure 1A**). The most caudal electrodes on each strip were physically connected to create a common reference. Retention rings with a dural replacement were installed at each burr hole to prevent bone growth and secure the EEG strips in place. The transceiver’s housing served as ground (Martinez-Ramirez, Slate et al. 2022). Video electrocorticography (ECoG) signals were recorded, and transmitter battery function/telemetry stability was assessed weekly to ensure signal quality. Pigs in Cohort 2 underwent an additional 3 surgeries for 2-photon imaging for a separate research objective.

**Figure 1.**
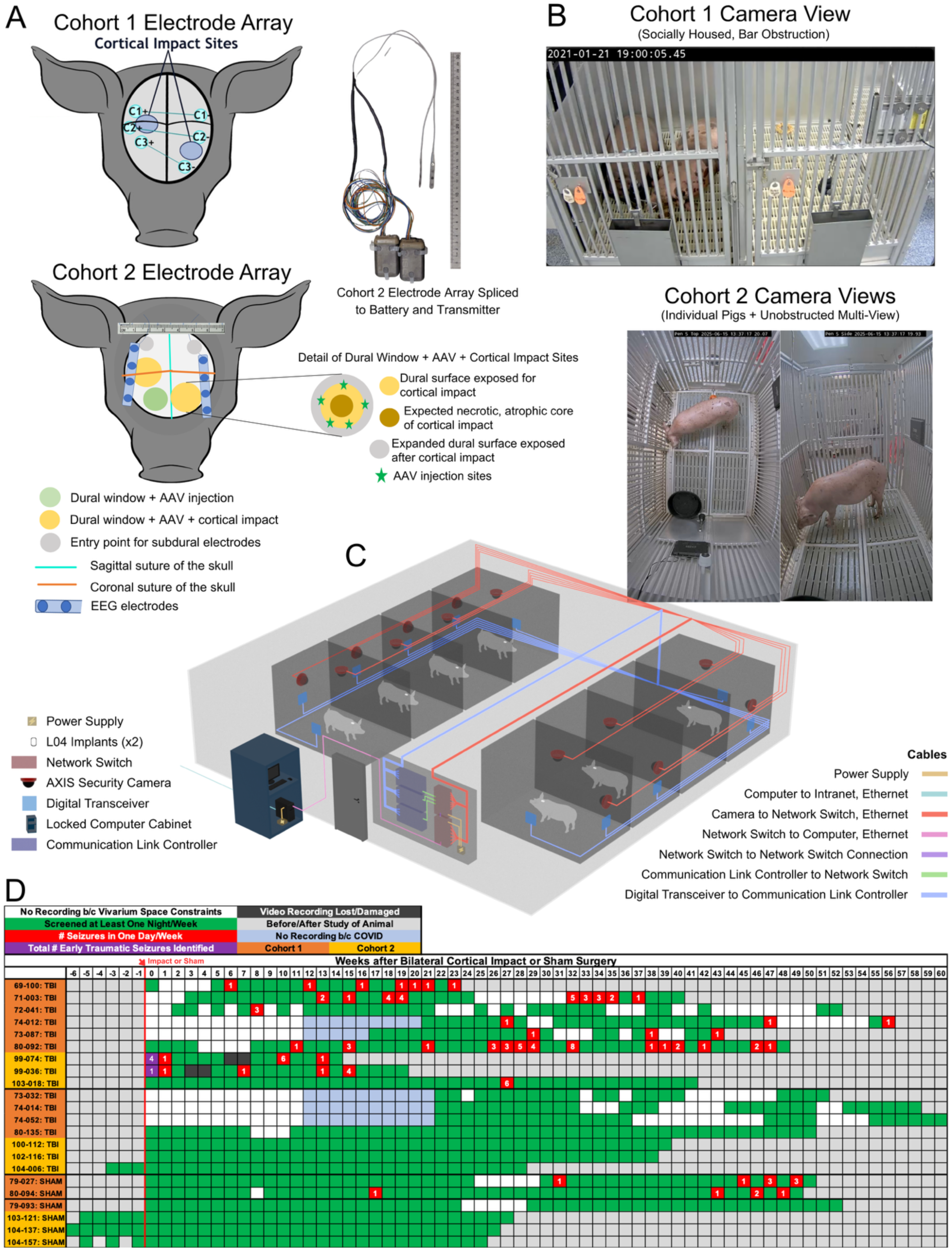
Evolution of long-term remote video electrocorticography (ECoG) and timing of the development of post-traumatic epilepsy (PTE) per pig. **A.** Location of bilateral cortical impact and ECoG electrode array for each cohort, including Adeno-Associated Virus (AAV) injection sites (subset of pigs). Electrodes were then spliced to the transceiver (L04) implanted in the neck. **B.** Comparison of improvement in recording techniques between cohorts 1 and 2. Cohort 1 had one camera view per recording, sometimes with multiple animals on one recording, possibly socially housed, and with an obstructed view due to pen bars. Cohort 2 had two cameras per pig and only one pig in view without any obstructing bars. **C.** Layout of the recording room, including recording equipment arrangement. **D.** Seizures documented through screening one day per week per pig across Cohorts 1 and 2. Subjects with **red** boxes had seizures with the number in the box as the number of seizures per day (24 hours) per week screened. Seizures can include multiple rounds of convulsions until normal behavior has resumed. The development of PTE was defined as two seizures greater than 7 days post-bilateral cortical impact or sham surgery. **Purple** indicates all early traumatic seizures observed ≤ 7 days after bilateral cortical impact (screened 24-7, not averaged per week/sum). **Green** indicates screened, but no seizure was observed. **Blue** indicates that subjects were not recorded due to being locked out of the lab due to COVID-19 restrictions. Gray indicates weeks before recording began or after the end of study for that pig, or weeks before impact that were not recorded. **Black** boxes indicate data was lost or failed to record properly.

During the Cohort 2 experiment, dexmedetomidine caused tissue necrosis in the ears of 4 pigs before that batch of drug was recalled and discontinued. The ear with the intravenous catheter was swollen and treated with 1% silver sulfadiazine ointment. No signs of infection were observed. Two pigs eventually underwent a pinnectomy to allow healing. Pig(s) from each treatment group were affected: 103-018 (TBI, PTE, pinnectomy), 103-121 (sham), 104-006 (TBI, no PTE), 102-116 (TBI, no PTE, pinnectomy). In none of these cases was the inner ear involved—only the pinna/skin of the ear flap was affected. No animals from Cohort 2 were excluded.

### Video ECoG Acquisition, Quality Control, and Storage

In both cohorts, pigs were video recorded during both day and night after cortical impact or sham surgery for up to 12 months (mean 40.5 weeks ± 13.5 SD). In Cohort 1, swine were recorded with a single video camera (AXIS M1145-L Network Cameras, Axis Communications; **Figure 1B**) and were socially housed on days when only video was recorded and individually housed for EEG and video recordings (Martinez-Ramirez, Slate et al. 2022). Recording every other week was attempted but was restricted by space limitations and the COVID-19 lockdown in 2020 (**Figure 1**). In Cohort 2, all pigs were individually housed throughout the study, and a subset of (N = 4) animals were video recorded prior to any surgery to assess normal behavior in naïve swine. In Cohort 2, synchronized video ECoG was collected 3-4 days per week for each animal, using two communication link transceivers to receive data from two transmitter implants per pig (**Figure 1C**). In one instance, a pig with TBI (no PTE) exhibited disruptive behavior, apparently bothered his neighbor, and was relocated to another pen within the room. Two additional naïve pigs were evaluated, age-matched to 120 and 180 days post-TBI or sham surgery, to assess normal behavior and check if any behavior described as peri-ictal and exclusive to PTE also occurred in naïve pigs.

Swine from Cohort 2 were recorded weekly for 36-72 hours using video data (Axis M3125-LVE Dome Camera) synced with ECoG via Ponemah software (DSI). This recording period included days prior to the first surgery to assess behavior in naïve pigs. Agent DVR software (iSpy) was used to continuously record 24 hours of video from 14 cameras. Videos were automatically saved to an institutional MAD3 network drive (massive array of disks; MassGeneral Brigham). The sampling rate of the activity channel was reduced to 1Hz to achieve a smaller file size that retained integrity. The video recordings were manually started and stopped weekly to maintain a manageable folder size and facilitate quality control, ensuring that all cameras were functioning correctly. A backup of the video & EEG data was stored on physical hard drives to prevent data loss. EEG was inspected periodically as an additional quality control measure. After the onset of PTE, pigs were continuously recorded until their battery were depleted. Battery life was expected to be 89 days per the manufacturer’s guidelines. However, discrepancies were noted between the expected and the reported battery remaining displayed by the software, requiring manual estimation of remaining battery life.

### Estrous cycle detection

Gilts in Cohort 2 were checked for signs of estrus 1-2 times daily as possible via applying pressure to their rear to detect standing estrus, as well as evaluation of vulvar swelling. In light-pigmented gilts, vulvar color was also evaluated with increased pink color. Estrous cycles were generally trackable through this process.

### Convulsion and Behavior Screening and Analyses

Seizures were defined by convulsions temporally associated with peri-ictal behaviors from the time when abnormal clusters of behavior were observed (**Table 1**) until the pig resumed normal behavior, including falling back asleep. A convulsion was defined as a distinct episode of high-intensity motor activity, typically consisting of involuntary, rapid, rhythmic muscle contractions and relaxations (clonic) and sustained contraction (tonic, **Table 1**). A seizure could be comprised of multiple episodes of convulsions. Convulsions were considered separate if there were ≥ 3 seconds of stillness between events. All potential convulsive events were reviewed in a group weekly to determine if they met the criteria in Table 1, and from videos of positive control convulsions and negative control convulsions (already reviewed and determined not to meet criteria). These behaviors, which might look odd and/or rhythmic but not convulsive, were then documented as negative criteria for what was not a convulsion (**Table 1**).

**Table 1.**
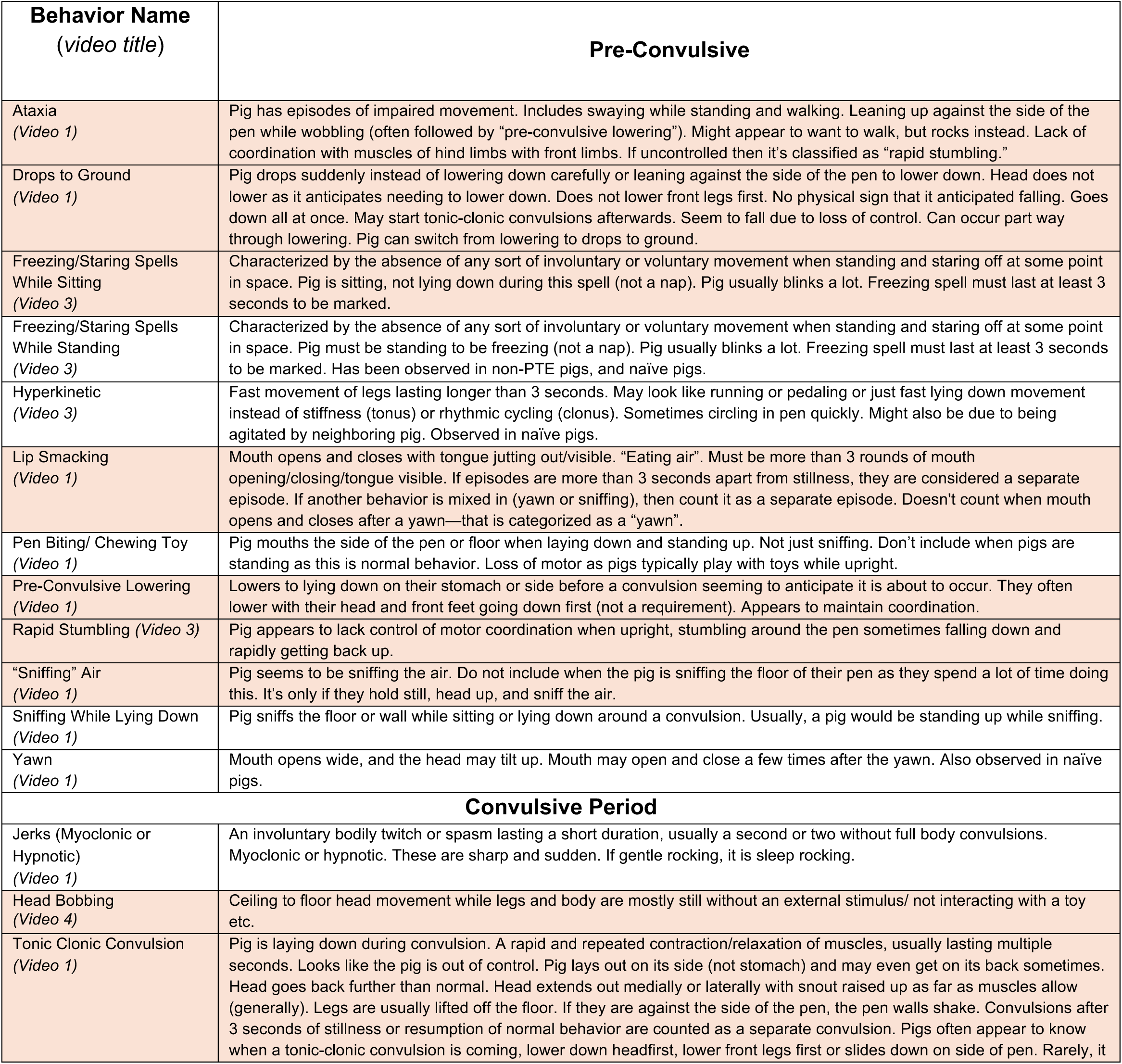
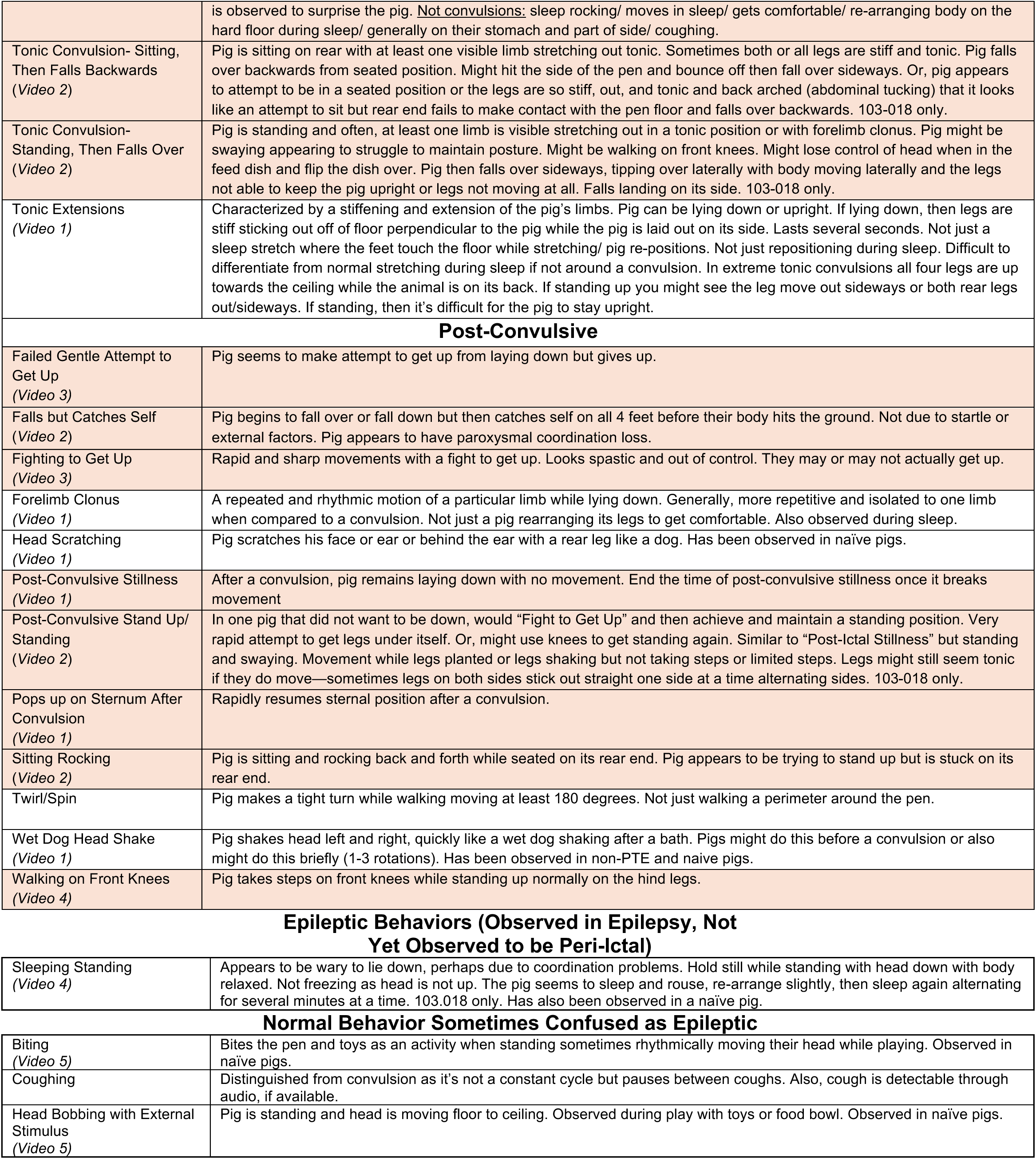
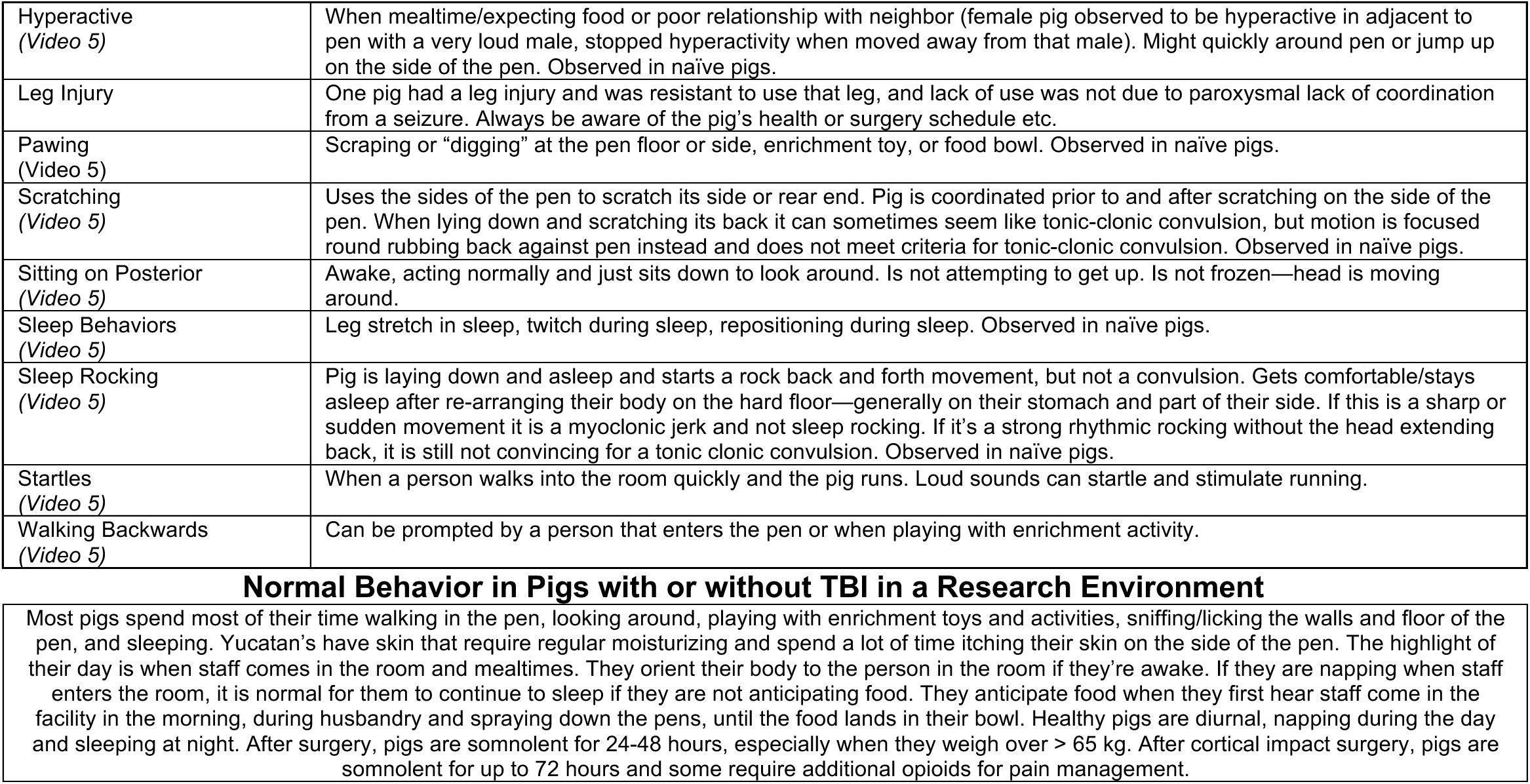
Behavior library of per-ictal vs. normal behavior. Peri-ictal behaviors were divided into pre-convulsive, convulsive period, and post-convulsive based on their general timing in documented events, averaged among all pigs with PTE (**Figure 2C**). Behaviors that were exclusive to epileptic pigs are marked in **orange**. Behaviors not in orange are observed around convulsions and were also observed in non-PTE pigs (alone, not clustered with convulsions). Epileptic behaviors observed in pigs with PTE but not yet observed to be peri-convulsive to date are noted. Rhythmic or odd behaviors that should not be considered epileptic are noted. Finally, how healthy pigs spend most of their time in a research environment is described. Video examples are cited per behavior. Behaviors are organized by domain, then alphabetized.

| Behavior Name<br>(video title) | Pre-Convulsive |
| --- | --- |
| Ataxia<br>(Video 1) | Pig has episodes of impaired movement. Includes swaying while standing and walking. Leaning up against the side of the pen while wobbling (often followed by “pre-convulsive lowering”). Might appear to want to walk, but rocks instead. Lack of coordination with muscles of hind limbs with front limbs. If uncontrolled then it’s classified as “rapid stumbling.” |
| Drops to Ground<br>(Video 1) | Pig drops suddenly instead of lowering down carefully or leaning against the side of the pen to lower down. Head does not lower as it anticipates needing to lower down. Does not lower front legs first. No physical sign that it anticipated falling. Goes down all at once. May start tonic-clonic convulsions afterwards. Seem to fall due to loss of control. Can occur part way through lowering. Pig can switch from lowering to drops to ground. |
| Freezing/Staring Spells While Sitting<br>(Video 3) | Characterized by the absence of any sort of involuntary or voluntary movement when standing and staring off at some point in space. Pig is sitting, not lying down during this spell (not a nap). Pig usually blinks a lot. Freezing spell must last at least 3 seconds to be marked. |
| Freezing/Staring Spells While Standing<br>(Video 3) | Characterized by the absence of any sort of involuntary or voluntary movement when standing and staring off at some point in space. Pig must be standing to be freezing (not a nap). Pig usually blinks a lot. Freezing spell must last at least 3 seconds to be marked. Has been observed in non-PTE pigs, and naïve pigs. |
| Hyperkinetic<br>(Video 3) | Fast movement of legs lasting longer than 3 seconds. May look like running or pedaling or just fast lying down movement instead of stiffness (tonus) or rhythmic cycling (clonus). Sometimes circling in pen quickly. Might also be due to being agitated by neighboring pig. Observed in naïve pigs. |
| Lip Smacking<br>(Video 1) | Mouth opens and closes with tongue jutting out/visible. “Eating air”. Must be more than 3 rounds of mouth opening/closing/tongue visible. If episodes are more than 3 seconds apart from stillness, they are considered a separate episode. If another behavior is mixed in (yawn or sniffing), then count it as a separate episode. Doesn’t count when mouth opens and closes after a yawn—that is categorized as a “yawn”. |
| Pen Biting/ Chewing Toy<br>(Video 1) | Pig mouths the side of the pen or floor when laying down and standing up. Not just sniffing. Don’t include when pigs are standing as this is normal behavior. Loss of motor as pigs typically play with toys while upright. |
| Pre-Convulsive Lowering<br>(Video 1) | Lowers to lying down on their stomach or side before a convulsion seeming to anticipate it is about to occur. They often lower with their head and front feet going down first (not a requirement). Appears to maintain coordination. |
| Rapid Stumbling (Video 3) | Pig appears to lack control of motor coordination when upright, stumbling around the pen sometimes falling down and rapidly getting back up. |
| “Sniffing” Air<br>(Video 1) | Pig seems to be sniffing the air. Do not include when the pig is sniffing the floor of their pen as they spend a lot of time doing this. It’s only if they hold still, head up, and sniff the air. |
| Sniffing While Lying Down<br>(Video 1) | Pig sniffs the floor or wall while sitting or lying down around a convulsion. Usually, a pig would be standing up while sniffing. |
| Yawn<br>(Video 1) | Mouth opens wide, and the head may tilt up. Mouth may open and close a few times after the yawn. Also observed in naïve pigs. |
| Convulsive Period |  |
| Jerks (Myoclonic or Hypnotic)<br>(Video 1) | An involuntary bodily twitch or spasm lasting a short duration, usually a second or two without full body convulsions. Myoclonic or hypnotic. These are sharp and sudden. If gentle rocking, it is sleep rocking. |
| Head Bobbing<br>(Video 4) | Ceiling to floor head movement while legs and body are mostly still without an external stimulus/ not interacting with a toy etc. |
| Tonic Clonic Convulsion<br>(Video 1) | Pig is laying down during convulsion. A rapid and repeated contraction/relaxation of muscles, usually lasting multiple seconds. Looks like the pig is out of control. Pig lays out on its side (not stomach) and may even get on its back sometimes. Head goes back further than normal. Head extends out medially or laterally with snout raised up as far as muscles allow (generally). Legs are usually lifted off the floor. If they are against the side of the pen, the pen walls shake. Convulsions after 3 seconds of stillness or resumption of normal behavior are counted as a separate convulsion. Pigs often appear to know when a tonic-clonic convulsion is coming, lower down headfirst, lower front legs first or slides down on side of pen. Rarely, it |
|  | is observed to surprise the pig. <u>Not convulsions</u>: sleep rocking/ moves in sleep/ gets comfortable/ re-arranging body on the hard floor during sleep/ generally on their stomach and part of side/ coughing. |
| Tonic Convulsion- Sitting, Then Falls Backwards<br>(Video 2) | Pig is sitting on rear with at least one visible limb stretching out tonic. Sometimes both or all legs are stiff and tonic. Pig falls over backwards from seated position. Might hit the side of the pen and bounce off then fall over sideways. Or, pig appears to attempt to be in a seated position or the legs are so stiff, out, and tonic and back arched (abdominal tucking) that it looks like an attempt to sit but rear end fails to make contact with the pen floor and falls over backwards. 103-018 only. |
| Tonic Convulsion- Standing, Then Falls Over<br>(Video 2) | Pig is standing and often, at least one limb is visible stretching out in a tonic position or with forelimb clonus. Pig might be swaying appearing to struggle to maintain posture. Might be walking on front knees. Might lose control of head when in the feed dish and flip the dish over. Pig then falls over sideways, tipping over laterally with body moving laterally and the legs not able to keep the pig upright or legs not moving at all. Falls landing on its side. 103-018 only. |
| Tonic Extensions<br>(Video 1) | Characterized by a stiffening and extension of the pig's limbs. Pig can be lying down or upright. If lying down, then legs are stiff sticking out off of floor perpendicular to the pig while the pig is laid out on its side. Lasts several seconds. Not just a sleep stretch where the feet touch the floor while stretching/ pig re-positions. Not just repositioning during sleep. Difficult to differentiate from normal stretching during sleep if not around a convulsion. In extreme tonic convulsions all four legs are up towards the ceiling while the animal is on its back. If standing up you might see the leg move out sideways or both rear legs out/sideways. If standing, then it's difficult for the pig to stay upright. |
| **Post-Convulsive** |  |
| Failed Gentle Attempt to Get Up<br>(Video 3) | Pig seems to make attempt to get up from laying down but gives up. |
| Falls but Catches Self<br>(Video 2) | Pig begins to fall over or fall down but then catches self on all 4 feet before their body hits the ground. Not due to startle or external factors. Pig appears to have paroxysmal coordination loss. |
| Fighting to Get Up<br>(Video 3) | Rapid and sharp movements with a fight to get up. Looks spastic and out of control. They may or may not actually get up. |
| Forelimb Clonus<br>(Video 1) | A repeated and rhythmic motion of a particular limb while lying down. Generally, more repetitive and isolated to one limb when compared to a convulsion. Not just a pig rearranging its legs to get comfortable. Also observed during sleep. |
| Head Scratching<br>(Video 1) | Pig scratches his face or ear or behind the ear with a rear leg like a dog. Has been observed in naïve pigs. |
| Post-Convulsive Stillness<br>(Video 1) | After a convulsion, pig remains laying down with no movement. End the time of post-convulsive stillness once it breaks movement |
| Post-Convulsive Stand Up/ Standing<br>(Video 2) | In one pig that did not want to be down, would "Fight to Get Up" and then achieve and maintain a standing position. Very rapid attempt to get legs under itself. Or, might use knees to get standing again. Similar to "Post-Ictal Stillness" but standing and swaying. Movement while legs planted or legs shaking but not taking steps or limited steps. Legs might still seem tonic if they do move—sometimes legs on both sides stick out straight one side at a time alternating sides. 103-018 only. |
| Pops up on Sternum After Convulsion<br>(Video 1) | Rapidly resumes sternal position after a convulsion. |
| Sitting Rocking<br>(Video 2) | Pig is sitting and rocking back and forth while seated on its rear end. Pig appears to be trying to stand up but is stuck on its rear end. |
| Twirl/Spin | Pig makes a tight turn while walking moving at least 180 degrees. Not just walking a perimeter around the pen. |
| Wet Dog Head Shake<br>(Video 1) | Pig shakes head left and right, quickly like a wet dog shaking after a bath. Pigs might do this before a convulsion or also might do this briefly (1-3 rotations). Has been observed in non-PTE and naïve pigs. |
| Walking on Front Knees<br>(Video 4) | Pig takes steps on front knees while standing up normally on the hind legs. |
| **Epileptic Behaviors (Observed in Epilepsy, Not Yet Observed to be Peri-Ictal)** |  |
| Sleeping Standing<br>(Video 4) | Appears to be wary to lie down, perhaps due to coordination problems. Hold still while standing with head down with body relaxed. Not freezing as head is not up. The pig seems to sleep and rouse, re-arrange slightly, then sleep again alternating for several minutes at a time. 103.018 only. Has also been observed in a naïve pig. |
| **Normal Behavior Sometimes Confused as Epileptic** |  |
| Biting<br>(Video 5) | Bites the pen and toys as an activity when standing sometimes rhythmically moving their head while playing. Observed in naïve pigs. |
| Coughing | Distinguished from convulsion as it's not a constant cycle but pauses between coughs. Also, cough is detectable through audio, if available. |
| Head Bobbing with External Stimulus<br>(Video 5) | Pig is standing and head is moving floor to ceiling. Observed during play with toys or food bowl. Observed in naïve pigs. |
| Hyperactive<br>(Video 5) | When mealtime/expecting food or poor relationship with neighbor (female pig observed to be hyperactive in adjacent to pen with a very loud male, stopped hyperactivity when moved away from that male). Might quickly around pen or jump up on the side of the pen. Observed in naïve pigs. |
| Leg Injury | One pig had a leg injury and was resistant to use that leg, and lack of use was not due to paroxysmal lack of coordination from a seizure. Always be aware of the pig's health or surgery schedule etc. |
| Pawing<br>(Video 5) | Scraping or "digging" at the pen floor or side, enrichment toy, or food bowl. Observed in naïve pigs. |
| Scratching<br>(Video 5) | Uses the sides of the pen to scratch its side or rear end. Pig is coordinated prior to and after scratching on the side of the pen. When lying down and scratching its back it can sometimes seem like tonic-clonic convulsion, but motion is focused round rubbing back against pen instead and does not meet criteria for tonic-clonic convulsion. Observed in naïve pigs. |
| Sitting on Posterior<br>(Video 5) | Awake, acting normally and just sits down to look around. Is not attempting to get up. Is not frozen—head is moving around. |
| Sleep Behaviors<br>(Video 5) | Leg stretch in sleep, twitch during sleep, repositioning during sleep. Observed in naïve pigs. |
| Sleep Rocking<br>(Video 5) | Pig is laying down and asleep and starts a rock back and forth movement, but not a convulsion. Gets comfortable/stays asleep after re-arranging their body on the hard floor—generally on their stomach and part of their side. If this is a sharp or sudden movement it is a myoclonic jerk and not sleep rocking. If it's a strong rhythmic rocking without the head extending back, it is still not convincing for a tonic clonic convulsion. Observed in naïve pigs. |
| Startles<br>(Video 5) | When a person walks into the room quickly and the pig runs. Loud sounds can startle and stimulate running. |
| Walking Backwards<br>(Video 5) | Can be prompted by a person that enters the pen or when playing with enrichment activity. |

Videos excluded from behavioral analysis included those with corrupted video files or visual obstructions. In Cohort 1, at least 1 24-hour period of video-EEG was screened per week, but was limited by space and the COVID-19 lockdown (**Figure 1D**). In Cohort 2, video-EEG was screened for 36-72 hours. In both cases, seizures are displayed per week in **Figure 1D**. Additional screening per convulsive pigs was often completed (**Supplemental Table 1**).

An accelerometer-based program was written in Python (Pretell and Costine-Bartell 2026) to extract high-movement periods from the European Data Format file for manual screening. Videos were concatenated from the same time period from these high-movement periods (plus 10 seconds before and after the high-force event) into video clips for manual behavioral analysis. The accelerometer detected 3 directions (x, y, z), reporting a total activity integral of the activity signal over the defined logging rate, normalized to a minute in gravity per direction. A range of at least –7 Gs to +7 Gs was provided, with a corresponding output from approximately 0 to 4095. A jerk value was calculated (JerkValue_i_ = C **⋅** √(X_i+1_−X_i_)^2^ + (Y_i+1_−Y_i_)^2^ + (Z_i+1_−Z_i_)^2^, where C = Sampling Rate **⋅** 3.5347. The default sampling rate for activity channels was 1 Hz and combined all 3 channels (m/s^2^). The program reported events over 400 m/s^2^ for the activity signal. While the events were identified via the program, the original video was reviewed to confirm whether each event was a convulsion. The detection of convulsive movements was validated across 12 convulsive events, in which 4 raters either manually screened 24 hours of video or screened the concatenated video. The detection rates were equivalent across methods, yielding 95% via manual analysis vs. 93% via the accelerometer tool. Regardless, many videos were analyzed manually when the EEG transceiver was not on.

Screened convulsions were reviewed and rejected or confirmed at a weekly meeting involving multiple screeners. Screeners were blinded to the pigs’ treatment, but it was often not possible to blind them to certain pigs, as specific behaviors were associated with specific epileptic pigs, and all pigs often had distinctive bodies and markings. Definitions of each peri-ictal or epileptic behavior were created to differentiate specific behaviors **(Table 1)**, and behaviors around convulsions were quantified. Behaviors common to all pigs that might be confused as epileptic (strange and/or rhythmic), including naïve pigs, are stated at the end of **Table 1**.

Temporal alignment of behavioral data allowed quantification of peri-ictal semiology. The last 10 seizures before the end of the study were analyzed per pig in Python (Pretell and Costine-Bartell 2026). If a pig had fewer than 10 seizures, all were analyzed. For each seizure, the onset time of the first convulsion was plotted as time = 0, and behavioral event onset was plotted relative to the convulsion and subsequent convulsions. Within a pig, data from multiple seizures were combined to assess the onset of each behavior type per convulsion.

### Video ECoG analysis for Electrographic Seizure

Although semiology alone can be used to identify and classify seizures (Fisher, Cross et al. 2017), we also evaluated EEG for electrographic seizures with a 60 Hz notch filter. Criteria for electrographic seizure were defined as spikes of ≥2 Hz for ≥10 seconds. Spikes were defined as signals with a deflection from baseline with an amplitude of at least 100 µV with a duration between 20-100 ms. The ECoG in **Figure 6** was additionally filtered by subtracting a low-pass-filtered copy of each channel from itself (i.e., channel_filtered_ = channel_original_ – lowpass_0.9Hz_ (channel_original_), with the low-pass filter set to 0.9Hz in the Neuroscore (DSI) software. Cohort 1 used a 3-channel bipolar montage (Martinez-Ramirez, Slate et al. 2022). Cohort 2 had a 6-electrode, flexible montage.

### Statistics

The number of behaviors around a convulsion was compared among convulsions in the same seizure with a one-way ANOVA followed by Tukey’s multiple comparison tests. A one-sided, paired t-test was used to determine if the number of behaviors per convulsion increased from early traumatic seizures to post-traumatic seizures.

## Results

### Development of Post-Traumatic Epilepsy

Following bilateral cortical impact, 9 of 16 pigs developed PTE (56%), defined as at least two spontaneous seizures occurring ≥ 7 days post-injury. This rate was the same in Cohort 1 and Cohort 2. The average onset of PTE occurred within 4.6 months (± 3.4 SD) post-injury (**Figure 1D**) when screened at a rate of 24 hours per week. The range of PTE onset when examining all data, including additional screening > 1 day/week, was 2 - 47 weeks post-TBI (**Supplemental Table 1**). The overall frequency of seizures from PTE was 0.38/day +/- 0.27 (SD; median 0.35, range 0.10-0.81) for the entirety of the recording period (day 8 post-TBI to the end of the experiment, **Figure 1D**). A total of 199 seizures were observed.

An estimated total of 26,338 hours of video (5,000 manual, 3,050 accelerometer tool for Cohort 1; Cohort 2: 13,152 accelerometer, 5,136 manual) were screened manually and with accelerometers to highlight rapid movement, followed by manual screening to identify normal behavior, peri-ictal behavior, and convulsions. Seizures were defined as convulsions temporally associated with peri-ictal behaviors from the onset of abnormal behavior until normal behavior resumed. A convulsion was defined as a distinct episode of high-intensity motor activity, typically consisting of involuntary, rapid, rhythmic muscle contractions and relaxations (clonic) and sustained contraction (tonic, **Table 1**). In a subset of pigs (N = 5, TBI; **Figure 1D**), video was recorded continuously for 24 hours a day during the first week after cortical impact or sham surgery to detect early traumatic seizures. In this subset, 2 of these pigs had early traumatic seizures and developed PTE. Interestingly, these two male pigs immediately went on to have a post-traumatic seizure the following weeks when screened more intensively than 24 hours/week.

The majority of convulsions were tonic-clonic (**Video 1, Table 1**), whereas only two pigs displayed tonic-only convulsions (**Video 2**, **Table 1**). One pig (103-018) exhibited tonic convulsions exclusively. A behavioral seizure was defined as the period of paroxysmal abnormal behaviors around consecutive convulsions. Across animals that developed PTE, seizures were most characterized by a focal motor onset that progressed to generalized seizure activity. Generalized onset events were uncommon and were observed in only a single case (**Video 2**). Sudden dropping to the ground prior to a convulsion was rare, as pigs appeared to learn the signs of an oncoming seizure and would lower purposefully (usually headfirst for tonic-clonic, or rear first for tonic) or lean against the side of the pen as they attempted to control lowering. Some pigs with PTE only had 2-3 seizures, while others had dozens of documented seizures (**Figure 1D**). Sham pigs from Cohort 1 that developed epilepsy had a different pattern, with months between the 1^st^ and 2^nd^ seizure (**Figure 1D**).

Several improvements of the electrode array and the video-EEG recording room were made between Cohorts 1 and 2. Two sham pigs from Cohort 1 developed epilepsy, whereas no shams from Cohort 2 developed epilepsy (**Figure 1D**). A key difference regarding the development of epilepsy in shams is the use of electrode strips rather than skull screws in Cohort 2 (**Figure 1A**). The electrode array was improved to accommodate additional channels with two EEG transceivers implanted. The additional channels facilitated remontaging, which off-the-shelf DSI implants do not. In Cohort 1, swine were transported to an outside warehousing facility due to a lack of space in the animal facility space, and recordings were obtained from a single camera that often captured multiple animals simultaneously and/or was partially obstructed by bars **(Figure 1B**). In Cohort 1, indoor cameras required protective covers (altered basketball display cases) that had to be closed during pen cleaning, while those in Cohort 2 were water-resistant outdoor security cameras. In Cohort 2, swine were continuously housed onsite in the recording room, with equipment expanded to record up to 7 swine simultaneously (**Figure 1B**). Swine from Cohort 2 were filmed continuously, 24 hours a day, 7 days a week, using 2 cameras per swine to capture both top and side views (**Figure 1C**). To prevent visual obstructions caused by bars in Cohort 1, cameras were relocated inside the pen (**Figure 1B**). Outdoor security cameras were already equipped with protective covers. These changes, along with the addition of video-ECoG monitoring, created conditions more favorable for capturing behavior and seizure events over extended periods.

### Peri-ictal Behavioral Library and Seizure Semiology

A total of 31 distinct behaviors that were motor, including apparent automatisms, and non-motor behaviors that occurred before, during, or after convulsions (peri-ictal behaviors) were identified via manual screenings (**Table 1, Videos 1-3, Figure 2**). The mean number of behaviors during a seizure was 7.3 ± 5.1 (SD; median 6, range 1-26). We divided these behaviors temporally connected to convulsions into 1) pre-convulsive, 2) convulsive period, and 3) post-convulsive (**Table 1**, **Figure 2**). Similar to people with epilepsy, these behaviors might be observed around a convulsion and also at other times with purposeful movement. For example, a person might have mouth movements and lose control of their saliva during a seizure, but mouth movements also occur during regular, controlled behavior. Therefore, the clustering of behaviors around convulsion is a key hallmark. Behaviors identified as exclusive in pigs with PTE are highlighted in orange in **Table 1**. Lip-smacking and sniffing air, potential automatisms, appeared to be specific to a subset of epileptic pigs. Epileptic/peri-ictal and non-epileptic/non-ictal behaviors were confirmed as being absent in naïve pigs prior to any anesthesia or surgery.

**Figure 2.**
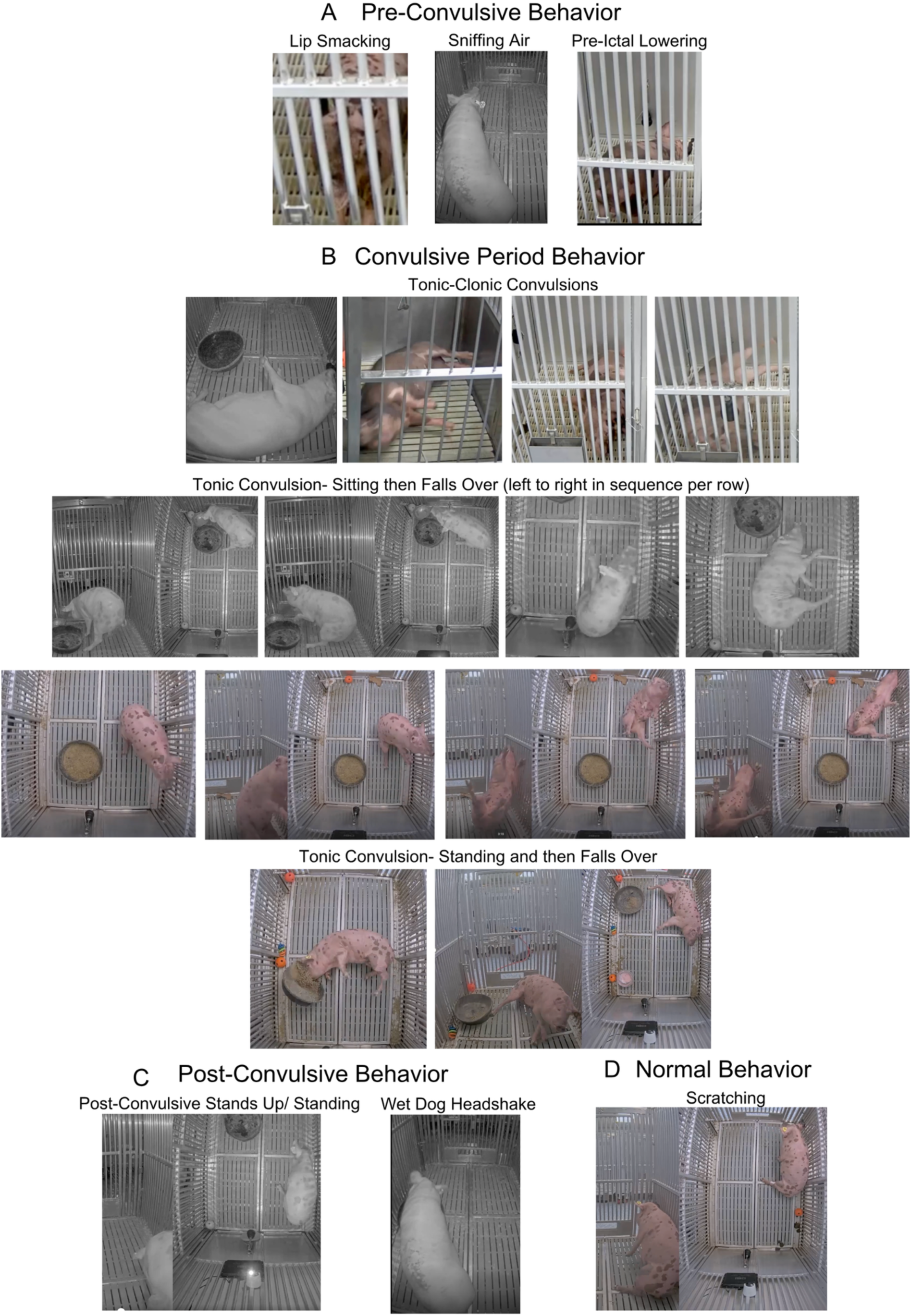
Pigs with post-traumatic epilepsy display a variety of unique behaviors. **A.** Pre-convulsive behavior displayed here is lip smacking, sniffing, and pre-ictal lowering. Pre-ictal lowering shows posterior still up and pig lowering on the front right shoulder first. **B**. When two camera angles are available, both are included if helpful, without space between views. Convulsions shown here are tonic-clonic, displaying front or rear legs and/or head extended out straight in 4 separate pigs. Two examples of Tonic Convulsion**—**Sitting, Then Falls Over, with still photos in order, left to right in sequence. The top row on the left shows the extreme abdominal tucking. Tonic Convulsion**—**standing then falling over displays two different events. **C**. Post-convulsive behavior includes post-convulsive stands up/ standing (note the tonic/extended left leg and abdominal tucking), and wet dog headshake. **D**. An example of normal behavior that might be confused as epileptic is a pig scratching against the pen.

We were careful to document several behaviors that were sometimes labeled as convulsions by screeners but did not meet the established criteria and are normal behaviors in swine. Staff training involved a review of epileptic and non-epileptic behaviors captured on video with concatenated EEG for certain examples. Care was taken to delineate stereotypical, non-pathological behaviors (sleep rocking, coughing) from epileptic ones. Biting, coughing, scratching, and sleep rocking, which are characterized by rhythmic movement (**Table 1**, **Video 5**) but are not epileptic. Behaviors that looked odd, but were not associated with seizures included hyperactivity, sitting on posterior, sleep behaviors, startle, and walking backwards (**Table 1, Video 5**). Other activity in the room must be taken into account. Pigs become hyperactive as they anticipate feeding, with increased EEG activity (Martinez-Ramirez, Slate et al. 2022). In one case, a pig in a pen next to a disruptive pig (100-112, TBI no PTE) was no longer hyperactive after relocation. Post-mortem analysis of the disruptive pig revealed a large area of tissue loss in the rostral, ventral portion of the left hemisphere in proximity to one of the cortical impact sites, not observed in other cases. Pigs might startle when someone enters the recording room. They walk backward while playing with enrichment toys or when a person enters the pen, making room for the person to enter. Finally, when healthy pigs sleep, they have periods of stillness intermixed with periods of substantial movement. Pigs, like people and dogs, make many movements while sleeping, including rocking, repositioning, stretching, and twitching (**Table 1, Video 5**).

### Quantitative Analysis of Peri-Ictal Behavior

Though the time spent in a convulsion was a few seconds, the pre-convulsion and post-convulsion behaviors could last for many minutes (maximum seizure length was 7.9 minutes); therefore, we screened for 3.5 minutes before and 4.5 minutes after each convulsion, or until regular behavior resumed (**Figure 3**). Seizures occurred while the pig was awake or after sleep. Sometimes, it was difficult to determine the end of post-ictal stillness because the pig would fall back asleep. Generally, pigs with PTE did not appear distressed by their uncontrolled behaviors (convulsions and associated behaviors), except for one pig (103-018), described below. Aside from loss of coordination and falling, none of the peri-convulsive or post-convulsive behaviors appeared to be severe. Convulsions are certainly dramatic (**Video 1**, **Video 2**), but no pig appeared to be apneic or to injure themselves, aside from some skin abrasions in Cohort 1 when the pen’s elevated floors were metal. No injuries were observed in Cohort 2 when the pen floors were switched to hard plastic. None died.

**Figure 3.**
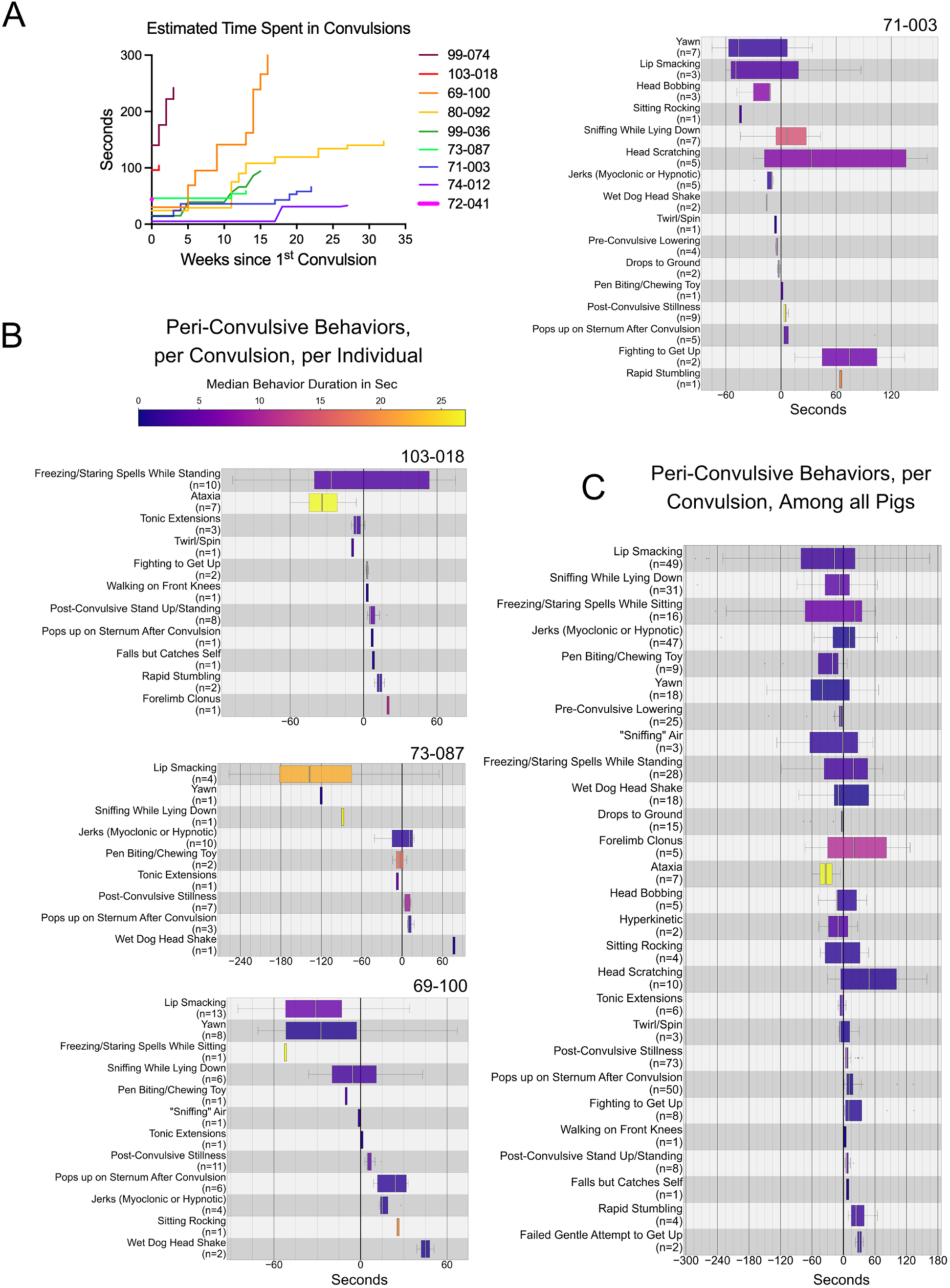
Pigs with post-traumatic epilepsy (PTE) demonstrate an array of peri-convulsive behaviors to varying degrees. **A**. Estimated time spent in convulsions per pig as PTE developed. Pigs had very brief convulsions, which were part of a wider array of behaviors within the entire seizure (defined as the entire period of abnormal behavior). **B**. Examples of per-ictal behavior per pig per convulsion. For each pig, its last 10 seizures (or all seizures if fewer than 10 were observed) were screened for defined behaviors (**Table 1**). The time difference between the onset of the behavior instance and its associated convulsion was calculated, plotted, and color-coded by the median duration of each behavior. The behavior and number of times observed are on the y-axis. The time between convulsion onset and peri-convulsive behavior onset is in seconds on the x-axis. The pig with the greatest array of peri-ictal semiology (71-003), the least (103-018), and in between (69-100) are demonstrated here. Box plots for the remaining pigs were plotted (**Supplemental** Figure 1). **C**. The compiled onset, duration, and temporal location of peri-convulsive behavior.

Each epileptic pig exhibited an individualized behavioral pattern, with some core elements displayed in most convulsions, indicating that seizure semiology was somewhat stereotyped within animals but heterogeneous across subjects (**Figure 2**). Some pigs had up to 22 identified peri-ictal behaviors, with 11 unique behaviors (72-041). Temporal alignment of behavioral onsets relative to convulsion onset (time = 0 seconds) of the last 10 seizures per pig with PTE (N = 87 seizures, N = 152 convulsions) revealed distinct peri-ictal patterns within and among pigs (**Figure 3B-C, Supplemental Figure 1**), with some behaviors being very common among all convulsions (lip smacking, post-ictal stillness in 69-100). Pre-convulsive behaviors included oral automatisms, such as lip smacking, yawning, and “cage/toy biting,” while lying down, as well as sniffing, including “sniffing while lying down” and “sniffing air,” which appeared like the pig could suddenly smell something (**Table 1**). An array of pre-convulsive behaviors also demonstrated coordination issues, including ataxia, “rapid stumbling,” and “drops to ground” or freezing (“freezing while standing” “freezing while sitting”). At other times, pigs appeared to purposely lower themselves down (“pre-ictal lowering”), sometimes using the pen side as support, as they seemed to anticipate the convulsion. Behaviors during the convulsive phase of the seizure included tonic-clonic convulsions, tonic convulsions with mixed in “tonic extensions” of legs, myoclonic jerks, and “head bobbing” (**Table 1**). The most consistent post-convulsive behaviors were “post-convulsive stillness,” where pigs would lie very still for typically 10-30 seconds after each convulsion followed by “pops up on sternum” (**Figure 3C**). Post-convulsion, pigs would sometimes make attempts to get back up that were not successful, including “failed gentle attempt to get up,” “fighting to get up,” and “sitting rocking.” Some pigs would scratch their head with their back leg (“head scratch”) or do a “wet dog head shake.” Though all these behaviors were observed around convulsions, not all of them were exclusive to pigs with PTE. For example, non-epileptic pigs were observed to “head scratch,” “yawn,” and “wet dog headshake” (**Table 1** orange boxes = exclusive to pigs with PTE vs. white boxes = not exclusive to pigs with PTE).

Specific differences in patterns of peri-ictal behavior, convulsions, and seizures among pigs were observed. 103-018 displayed a combination of completely unique peri-ictal behaviors, as well as peri-ictal behaviors common to other epileptic pigs (**Figure 3B**, **Table 1**). Prior to developing PTE, 103-018 had episodes of falling, jerks while lying down, slow lowering while on her front knees, and episodic loss of balance. At 27 weeks post-TBI, she started displaying persistent ataxia and had her first convulsive seizure later the same day, and then her second seizure the day after that. While other pigs seemed to want to lie down before a convulsion, 103-018 seemed to resist lying down and even appeared to sleep while standing for prolonged periods of time (**Video 4**). The maximum length of sleep standing was 36 hours at week 27 post-TBI, the day after she developed persistent ataxia. After the development of PTE, she spent much of her day standing and freezing and standing and sniffing more than other pigs. She was the only pig to have exclusively tonic convulsions (one other pig had a mix of tonic and tonic-clonic convulsions); we identified two types: “Tonic Convulsions—Sitting, Then Falls Backward” and “Tonic Convulsions—Standing, Then Falls Over” (**Table 1**, **Figure 3B, Video 2, 3**). In both cases, she completely lost coordination and would fall over. Instead of “Post-Convulsive Stillness”, lying still, and then “Pops up on Sternum After Convulsion” like other pigs, she would get up suddenly on all four legs and then stand relatively still while swaying. These seemed comparable to “Post-Convulsive Stillness”, but she appeared to work hard to stand, so we combined these behaviors into one, “Post-Convulsive Stand Up/ Standing”. In the final month of observation, no convulsions or epileptic behavior were observed while the varying degrees of ataxia continued. 74-012 was notable as he displayed an array of peri-ictal behavior (**Figure 3B**) for several months before the first seizure (at 6.75 months) and second seizure at 11.75 months post-impact (**Figure 1D**). This pig exhibited a wide array of peri-ictal behaviors prior to PTE onset, with a limited number of observed convulsions.

Each seizure often had multiple rounds of convulsions during its duration (**Figure 4**), with the longest seizure lasting 7.9 minutes (99.074). If a seizure had more than one convulsion, the number of distinct behaviors was greatest around the first convulsion, with fewer behaviors associated with subsequent convulsions (**Figure 4A**, **Supplementary Figure 2**). The greatest number of rounds of convulsions within a seizure observed was 7 (**Figure 4B**, 73-087). Most pigs displayed the array of peri-ictal behavior around the first convulsion and not the subsequent convulsions, with a subset of pigs displaying a significant number of peri-ictal behaviors around the subsequent convulsions (69-100, 72-041, 73-087, **Figure 4**; 72-041, **Supplemental Figure 2**).

**Figure 4.**
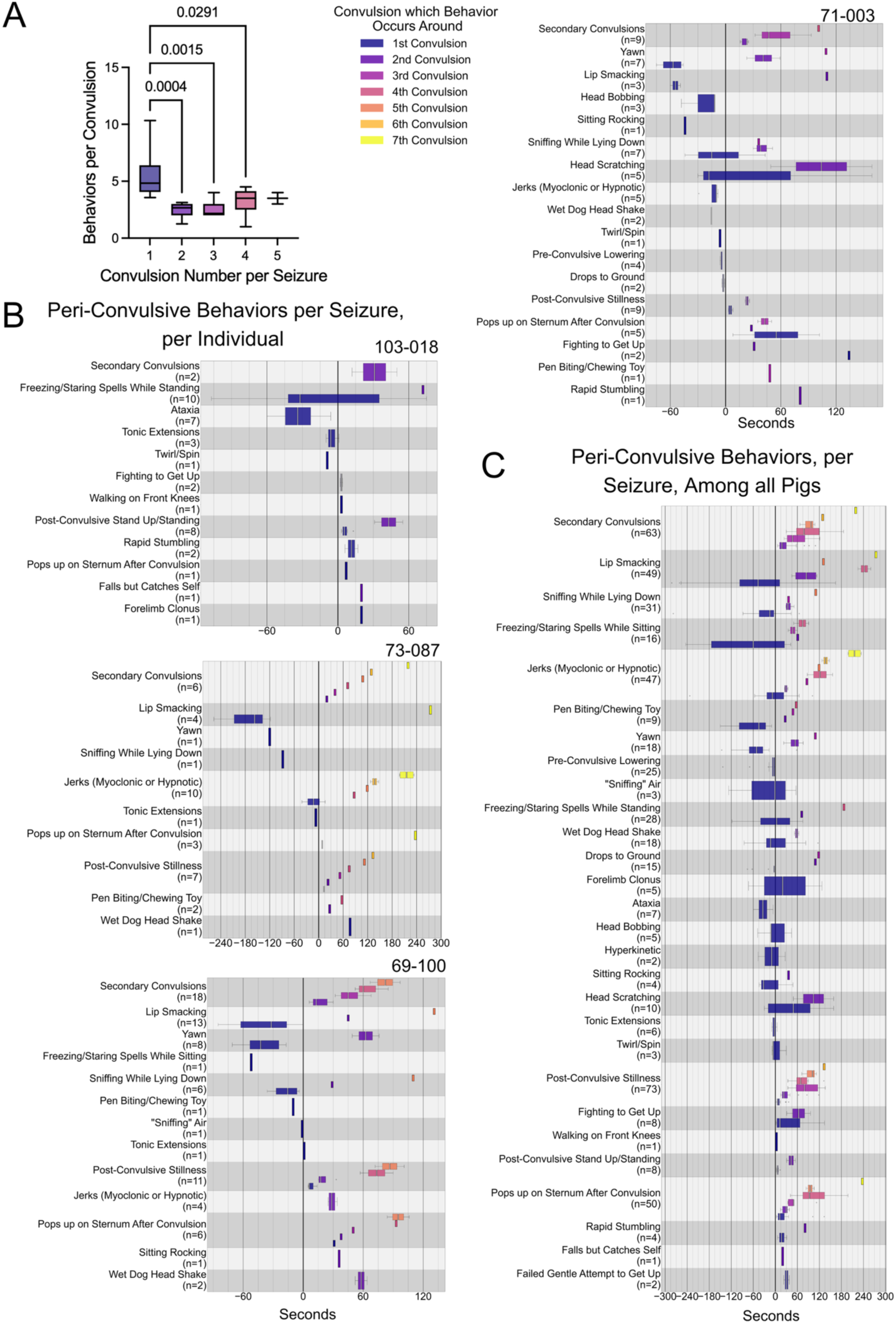
Semiology of per-ictal behaviors around convulsions within the same seizure. **A.** The number of behaviors was greatest around the first convulsion of a seizure vs. successive convulsions (one-way ANOVA) **B**. Semiology of seizures for an individual pig, with the onset of the first convulsion set as zero on the x-axis and the timing of each behavior (y axis) around each consecutive convulsion plotted per color (e.g. convulsion 1 = navy etc.). During a seizure, some behaviors continued to be displayed in later convulsions. **C.** The compiled onset and duration of peri-ictal behaviors among all pigs during seizures. The longest seizure was 7.9 minutes.

Of females in Cohort 2, where the estrous cycle was tracked, only one developed PTE (103-018). Her estrous cycle was detectable and regular for 7 cycles. The estrous cycle before the development of PTE was short (9 days), and then she appeared to be persistently in standing estrus prior to and during the development of PTE. She might have had a follicular cyst. After the development of PTE, her seizures were clustered, her estrous cycles were irregular, and she might have had additional follicular cysts. Stress can cause follicular cysts. Alternatively, the detection of standing estrus may have been obscured by persistent ataxia.

As Cohort 1 was a pilot project, we had a limited number of pigs for which we performed video-EEG prior in the week following cortical impact (n = 5, both cohorts). Two of these pigs displayed early traumatic seizures (ETS) that were similar, yet distinct from the seizures they developed later (**Figure 5**). Some elements remained the same from ETS to PTE, but some peri-ictal behaviors present in PTE were absent in ETS (**Figure 5D**, red boxes). The average number of per-ictal behaviors tended to increase from ETS to PTE (2.2 vs. 5.4; **Figure 5D**; P = 0.11).

**Figure 5.**
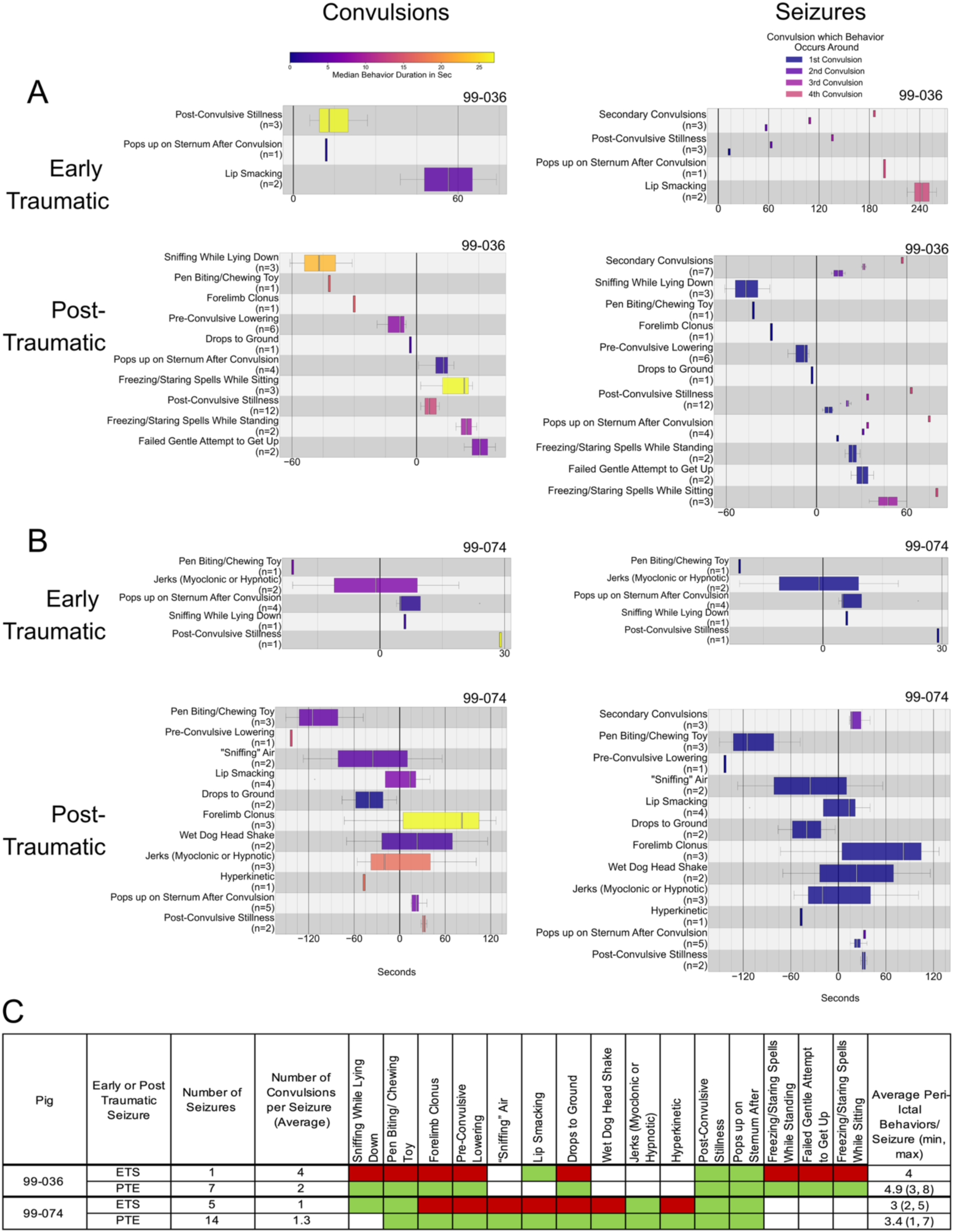
Inventory of peri-ictal behaviors in pigs with both early traumatic seizures and post-traumatic seizures. **A.** Early traumatic convulsions (left) and early traumatic seizures (right) in pig 99-036. **B.** Early traumatic convulsions (left) and early traumatic seizures (right) in pig 99-074. **C.** Behaviors per seizure increased from early traumatic seizures (ETS) to post-traumatic seizures (PTE). **Green boxes** mark behaviors present in PTE seizures. **Red boxes** in ETS indicate behavior present in PTE seizures but not in ETS seizures per pig. ETS = 0-7 days after bilateral cortical impact; PTE, > 8 days after bilateral cortical impact.

**Figure 6.**
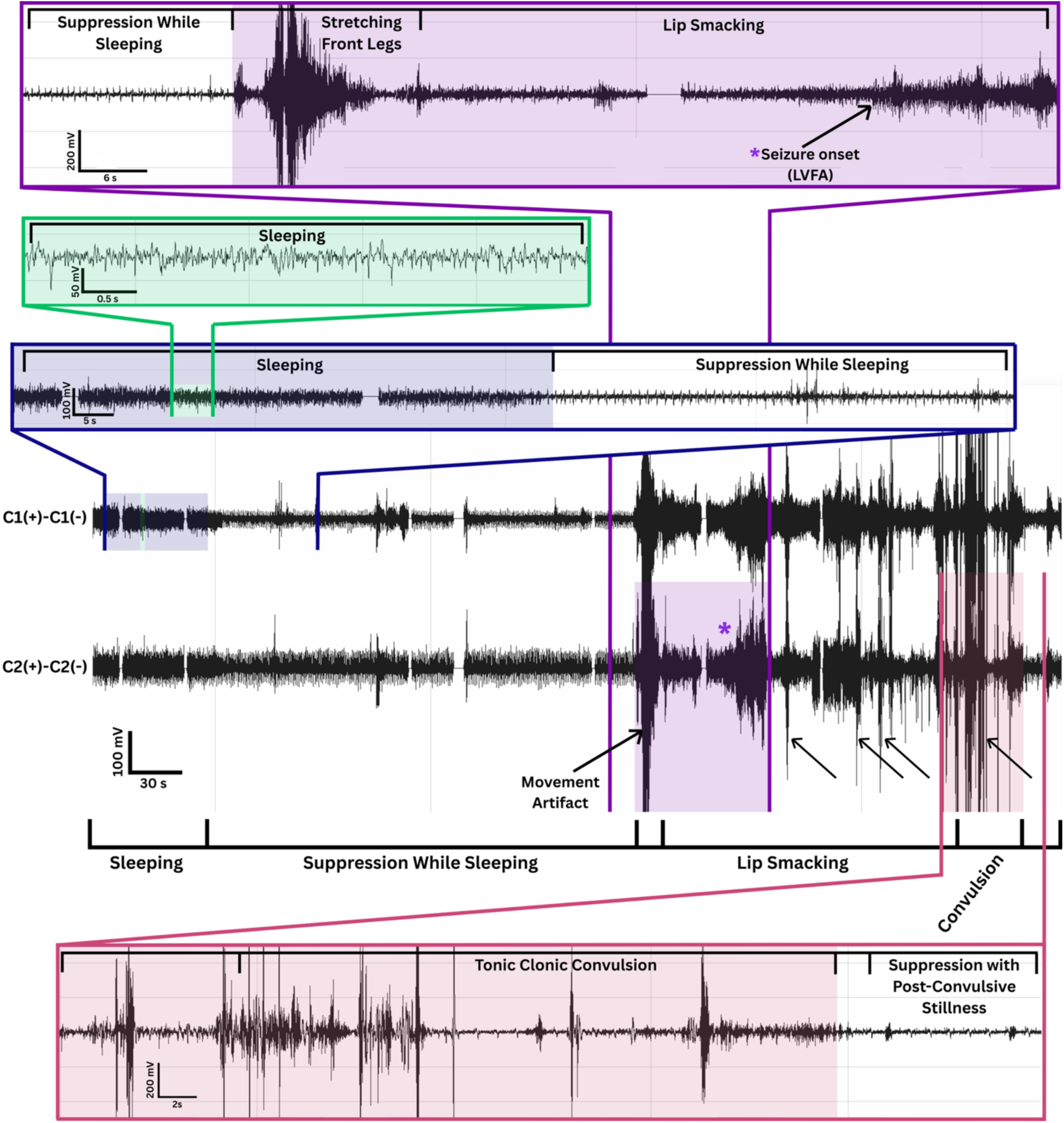
Example of electrographic seizure occurring during epileptic behaviors. The overview of the electrocorticography (ECoG) is the center 2 channels with time expansions of ECoG above and below in colored boxes marking ECoG activity at the time of peri-ictal behaviors. Electrographic seizure onset was characterized by suppression, followed by low-voltage fast activity (*; purple box), both occurring prior to the onset of pre-convulsive semiology, and then was followed by post-seizure suppression. Semiology began with automatisms such as lip smacking that evolved into a tonic-clonic convulsion, followed by post-convulsive stillness. During lip smacking, the ECoG shows that amplitude increases despite the movement remaining constant. **\***Low voltage fast activity. **Arrows**: movement artifact prior to seizure (leg stretch) and during the seizure (cage biting, tonic-clonic convulsion).

Some behavioral expressions of convulsions were accompanied by video electrographic activity (**Figure 6**), whereas other electrographic activity was obscured by movement artifact. In one example (**Figure 6**), the seizure began with low-voltage fast activity (Frauscher, Mansilla et al. 2024) followed by an increased amplitude coinciding with semiologic evolution (**Figure 6**).

## Discussion

To our knowledge, this is the first characterization of PTE in swine. Our 56% rate of PTE in each cohort is surprisingly high, as only multi-lesion, multi-insult (MuLMI) subarachnoid/subdural hemorrhage and/or contusion and/or penetrating with or without periods of apnea and early traumatic seizures in humans generate this high a rate of PTE (Raymont, Salazar et al. 2010). Patients with this load of MuLMI generally require ventilation and intensive care support and have a low Glasgow Coma Score. We do not hypothesize that swine have a special resistance to MuLMI as we can certainly induce MuLMI in pigs, resulting in lower neurological scores and requiring sedation and mechanical ventilation during the ictal phase (Costine-Bartell, Price et al. 2021). In the present bilateral cortical impact model, subjects fared well clinically in the acute phase post-injury, exhibiting somnolence in the first 24-48 hours but remaining ambulatory and able to eat. The low clinical burden of this TBI model might be due to the bilateral administration of the cortical impact in these specific rostral regions and offset locations of impact. A significant advantage of this model is the pathologically and chronically clinically significant injury (as far as PTE), without significant neurologic deficits in the acute phase, and even minimal early traumatic seizures. Again, this lends evidence to the disconnect between early traumatic seizures and post-traumatic epilepsy (Young, Rapp et al. 1983, Temkin, Dimken et al. 1990, Young, Okada et al. 2004). Fluid percussion injury in rats resulted in a similar rate of PTE at 50%, and the time for epileptogenesis after PTE depends on the level of monitoring, and has recently been reported as 22% (Kharatishvili, Nissinen et al. 2006, Ndode-Ekane, Ali et al. 2024, Andrade and Pitkänen 2026). It is well established that rats have a seizure rate of 0.2/day after lateral fluid percussion. We observed a rate of 0.38/day, but it was highly variable among pigs.

We hypothesize that the length of the latent period and the degree of variability are positively correlated with brain size among species. When expressed in terms of the unit of measurement used in rodents (days), we demonstrate a highly variable, prolonged average latent period of 198 days, with a range of 49-329 days. This is longer than in mice, which also display high variability from 5 to 126 days (Shandra, Mahmutovic et al. 2026), and comparable to the length in rats, which have a fairly uniform onset around 200 days (Andrade and Pitkänen 2026). Differences across these studies include the method of TBI: bilateral cortical impact in this study (52% developed PTE), lateral fluid percussion in rats (50%)(Kharatishvili, Nissinen et al. 2006, Pitkänen, Bolkvadze et al. 2011), and mild, repetitive weight drop in mice (25-44% developed PTE)(Shandra, Winemiller et al. 2019, Shandra, Mahmutovic et al. 2026), as well as the method and frequency of observation. Common among species, including here, is that implantation of EEG monitoring equipment can induce seizures, which are characteristically different than PTE (Andrade and Pitkänen 2026). In Cohort 1, one sham that developed epilepsy appears to have had a congenital deformity that also causes epilepsy in humans, and the other appeared related to the skull screws, but both shams with epilepsy had fewer seizures and a prolonged time between the first and second observable seizure. We seem to have avoided this problem in Cohort 2 by not using skull screws but epidural electrode strips instead. Among all species, some individuals displayed early traumatic seizures that were distinct in semiology from PTE.

A subset of pigs demonstrated seizure clustering similar to that described in rats (and humans), but outbred Yucatan pigs are likely more variable than rats, and other pigs did not show seizure clustering. In the female with PTE, where the estrous cycle was tracked, several estrous cycles were regular until the cycle prior to PTE development, which was short; after that, she was in prolonged standing estrus and might have had a follicular cyst. After the development of PTE, her cycles became irregular. Certainly, the estrous cycle can influence seizure generation, but stress and/or epilepsy might also disrupt normal estrous cycles (Scharfman and MacLusky 2014). Once this gilt developed PTE, she had prolonged periods of “sleep standing” and ataxia, which might have been stressful.

To our knowledge, this is also the first detailed description of seizure semiology in swine (Martinez-Ramirez, Slate et al. 2022). The array of behaviors we describe here in swine with PTE is extensive, similar to that of humans, cats, and dogs, and different from the limited behavioral repertoire of mice. Seizures began focally and became generalized, resulting in tonic-clonic or clonic convulsions. Pigs had seizures that lasted several minutes, with a maximum of 7.9 minutes. This is five times as long as spontaneous seizures in the mouse (Reddy, Golub et al. 2025) and three times as long as the rat (Andrade and Pitkänen 2026) models of PTE (90 seconds in mice, 150 seconds in rats). Additionally, pigs exhibited an array of per-convulsive behaviors, and a seizure can consist of multiple rounds of convulsions, with a maximum of 7 observed. Perhaps seizure length and the array of peri-ictal behavior are also positively correlated with brain size.

We describe a behavioral library of 31 behaviors associated with PTE, which could be expanded to about 100 if subcategorized by direction or per limb, as carefully described in rats (Andrade and Pitkänen 2026). Core behaviors common among species include freezing, limb clonus, sniffing, jerks, and head bobbing. Additionally, we describe normal behavior that might be mistaken for epilepsy in swine, as well as normal behavior in a research facility context. Examples of behaviors observed in rats after PTE that we did not observe include body curling, agitation, exploration with sniffing (close to our “sniffing the air”), whisker tremors, slow wandering, exploration, and rearing (Andrade and Pitkänen 2026). The majority of pigs with PTE described had tonic-clonic convulsions (89%, one pig had tonic-only), but these were rare (6-17%, “clonic body jerks”) in rats with PTE (Andrade and Pitkänen 2026). Pigs might have a wider natural behavior repertoire than rats, so that slow wandering and exploration are not distinctive (common in healthy pigs) or require machine learning for detection. Examples of behaviors in mouse models of convulsant-induced epilepsy that we did not observe in pigs with PTE included clonic convulsions while sitting, tonic-clonic convulsions while lying on the stomach (pigs lie on their side), wild jumping, apnea, and death (Van Erum, Van Dam et al. 2019). Pigs did not exhibit some behaviors observed in epileptic dogs, such as panting, anxiety, howling, whining, and jaw clonus (Berendt, Gredal et al. 2004). This suggests that the ictal repertoire is rooted in the species’ natural behavior, per their anatomy and agility. Here, we use the ILAE guidelines for describing the seizure: motor or non-motor onset followed by generalization (Fisher, Cross et al. 2017) and timing of the behavior to before or after convulsion, and not the Racine scale, which is for assessing seizure severity (Racine 1972, Van Erum, Van Dam et al. 2019). We agree with Andrade and Pitkänen that seizures from organic etiologies might be less severe than induced seizures, and the Racine scale, developed using electrical current or chemo-convulsant, is not applicable to PTE, but a behavioral library is more appropriate (Andrade and Pitkänen 2026). Seizures that develop naturally appear to become more complex over time, but their severity and lethality are lower than in chemical-induced models. Complexity is not equivalent to severity.

Pigs displayed a range of behaviors similar to those of cats, dogs, and humans, but absent in rodent models of epilepsy (Beniczky, Tatum et al. 2022). This included an array of complex facial automatisms with the tongue jutting out (“lip smacking”) and circling (“twirl/spin”), similar to those seen in cats with naturally occurring encephalitis (Binks, Crawford et al. 2025). Swaying and crawling observed in dogs was encompassed in “ataxia” in the pigs here (Berendt, Gredal et al. 2004). While we could not survey the pigs on their sensory, affective, cognitive, and autonomic symptoms (Beniczky, Tatum et al. 2022), we did detect automatisms (“lip smacking”, “sniffing air”), and many pigs seemed to be aware that convulsions were going to develop, few pigs seemed to fall down, but purposefully lowered, attempting to reach the floor of the pen prior to convulsion.

Small-scale phenotypes such as mydriasis (cats), salivation (rats and cats), chewing (rats), and ear movements (rats and dogs), eye blinking (rats), fast breathing (rats), scanning the environment with their head (rats), lacrimal secretion (dogs), and tail extension (rats) might have been observable in pigs but not possible at the video resolution here used to record for several months. Additionally, we did not record audio and therefore, could not observe vocalizations described in mice, rats, cats, and dogs. Future work might focus on known epileptic pigs more intensively with additional cameras, audio recording, and determining the level of consciousness by stimulation during seizure.

Identifying epileptic behavior in pigs required special considerations distinct from those for singly housed rodents or dogs. Those not knowledgeable and experienced with livestock species should be careful not to confuse normal behavior as abnormal, using **Table 1** to distinguish one from another. We did not include agitation as a potential behavior connected to PTE, as pigs were in pens open to neighbors, and social conflicts arose between some subjects with TBI (with or without PTE). Pigs are very social. We recommend that they be individually penned, but still have visual, physical, and odor cues from neighbors for animal welfare reasons. Therefore, social interactions must be considered, with staff spending time in the room with the pigs to understand social dynamics, rather than only watching videos disconnected from the room’s context. It is not advised to group/house pigs in the same pen for the detection of epilepsy, because when pigs are in the same pen, the amount of behavior tied to social interactions increases, as does visual obstructions. If pigs sleep in the same pen, the movement of one pig is transmitted to the other, and the source of the movement is obfuscated. Additionally, group housing increase the risk of misidentifying pigs and interferes with autonomous characterization of behavior with machine learning. The most common behavior mis-assigned by raters was sleep rocking and sleep behaviors; healthy, naïve pigs have a surprisingly large amount of rhythmic movement during sleep. Pigs without PTE exhibit a wide range of movements while sleeping, including rearranging, stretching, and sleep rocking, in which the pig moves side to side in a rhythmic fashion. Training with video examples and written descriptions, with regular group review meetings, was needed to distinguish sleep from convulsions.

### Limitations

The study population comprises two cohorts: Cohort 1 was a pilot study and was not randomized, and both cohorts are small. Seizure incidence might be underestimated in this set of pigs because the pigs were not recorded in Cohort 1 for long periods due to space limitations and the COVID lockdown. Due to the long latent period in this species, pigs might have developed PTE after the recording period. However, in those that did eventually develop PTE had an array of clustering potentially epileptic behavior prior to convulsion, whereas other pigs had no potentially epileptic behavior. Additionally, pigs might have focal seizures that do not generalize and were missed by evaluating for convulsions only. More specifically, epileptic behaviors might exist with higher-resolution video. Not every seizure was validated with EEG. This study evaluated the semiology of seizure in a single type of epilepsy associated with a single TBI mechanism. While there might be differences in the semiology of other types of epilepsy (i.e., genetic epilepsies), many are likely to overlap, and the behavior of healthy pigs in a research setting should be consistent.

### Future Studies

The advantage of this model is a distinct latent period similar to that of humans, but it is also a disadvantage from a logistical standpoint, as the study might take a very long time to test therapeutics, etc. To harness the wide repertoire of behaviors during epileptogenesis, we are using machine learning to examine the behavior cataloged here and determine whether the frequency of these behaviors at times not temporally connected to convulsions is higher in epileptic vs. non-epileptic pigs, and whether it is possible to predict PTE. Sleep disruption and other epileptic semiologies presented here, as well as the patterns and clustering (paroxysms) of behaviors, might be detectable prior to PTE onset with self-training visual neural networks.

## Conclusion

In this first model of PTE or any type of epilepsy in swine, bilateral cortical impact resulted in PTE in 56% of subjects, with a prolonged, variable latent period and tonic-clonic and tonic convulsions, along with an extensive array of stereotypical, peri-ictal behaviors more similar to those of humans and other large animals than mice. Long-term, synchronized video-ECoG and behavioral annotations revealed stereotypical peri-ictal behavioral patterns that vary across individuals. This behavioral library will likely be applicable to other models of PTE and other causes of epilepsy modeled in swine. Investigations of epileptic behaviors will build upon this work with the aim of predicting which animals will develop PTE, improving our understanding of PTE, and aiding in the development and testing of targeted interventions to prevent PTE.

## Acknowledgements

We thank the Knight Surgical Research Laboratory for their assistance with anesthesia and surgical procedures. We thank Dr. Ann-Christine Duhaime for performing surgeries. We thank the team at the Center for Comparative Medicine for their animal husbandry, enrichment activities, and the excellent care of our animals. Special thanks go to Kimberly DeGrenier, Titilayo Lamiti, and Dr. Corina Beale for their logistical support in making this happen, and to Drs. Andrea Slate and Bernard Varian. We thank the many MGH brain trauma interns and technicians over the years for their assistance with animal work.

## Data availability

Tabular data with Common Data Elements labeling is available on Open Data Commons for Traumatic Brain Injury (Pretell and Costine-Bartell 2026). The code for the accelerometer tool and the code to create box plots of behavior are available on our lab’s GitHub (GitHub 2026). Video and ECoG are too large to be archived in data repositories and are available via reasonable request.

## Declaration of generative AI and AI-assisted technologies in the manuscript preparation process

The authors used Claude (Anthropic) for debugging and writing a small portion of code snippets. After using this tool, the authors reviewed, tested, and edited all code and take full responsibility for the content of the published article.

## Titles of Videos

**Video 1. Tonic-clonic convulsions and associated behaviors.**

**Video 2. Tonic convulsions and associated behaviors.**

**Video 3. Other peri-ictal behaviors.**

**Video 4. Behaviors specific to epilepsy, but not around a convulsion.**

**Video 5. Normal behaviors that should not be considered epileptic.**

