## Supplemental Materials for "Individualized and stereotypical seizure semiology in a porcine model of post-traumatic epilepsy"

**Supplemental Table 1. Data sets included for each analysis.**

| Characteristic | Includes shams with epilepsy? | Includes ETS? | 24 hrs/week both cohorts | Last 10 seizures of each PTE pig for box plots | All available data | Result |
|---|---|---|---|---|---|---|
| Seizure frequency number (text data) | No | No | 24 hrs/week (to avoid over sampling from some pigs) | N/A | No | 0.38/day +/- 0.27 (SD; median 0.35, range 0.10-0.81) for the entirety of the recording period |
| PTE onset average (text data) | No | No | 24 hrs/week (to avoid over sampling from some pigs) | N/A | No | 4.6 months (± 3.4 SD) |
| PTE onset range (text data) | No | No | No | N/A | All the data | ranging from 2 to 47 weeks post-TBI |
| Figure 1D | Yes- labeled as such | Yes- labeled as such | 24 hrs/week (to avoid over sampling from some pigs) | N/A | No | Figure 1D |
| Average Seizure Length (text data) | No | No | N/A | Yes | No | 109.5 seconds |
| Max seizure length | No | No | N/A | Yes | No | 7.9 minutes |
| Behaviors around convulsions boxplots | Yes, labeled as such | Yes | N/A | Yes | No | Figures 3 and 4 |
| Total number of seizures identified: both ETS and PTE and shams | Yes | Yes | No | No | Yes | N = 199 seizures |
| Total number of seizures analyzed with box plots | Yes | Yes | No | Yes | No | N = 87 seizures |
| Total number of convulsions analyzed in box plots | Yes | Yes | No | Yes | No | N = 152 convulsions |
| Time spent in convulsions panel (Figure 3A) | Yes | No | No | N/A | No | Figure 3A |

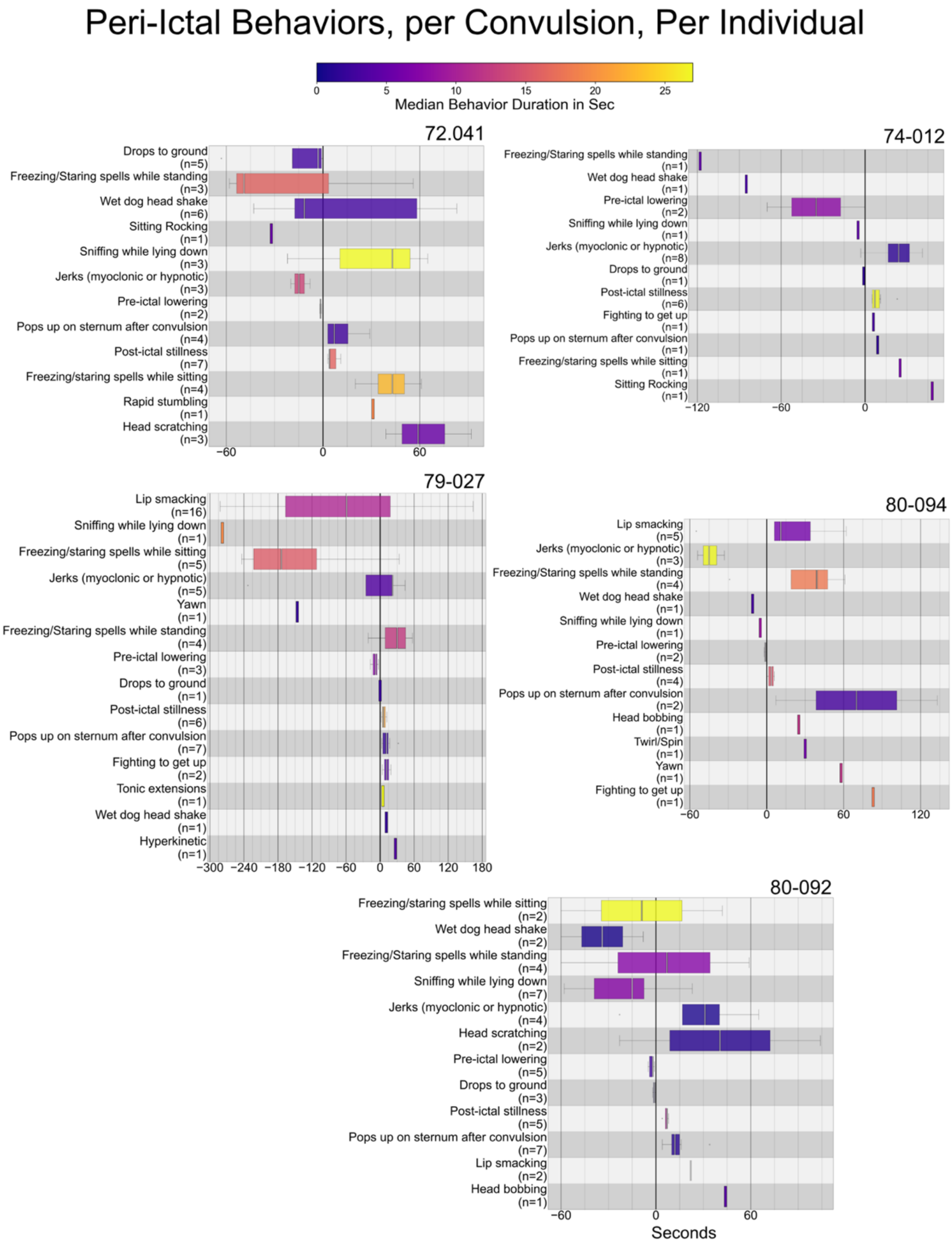


**Supplemental Figure 1. Peri-ictal semiology per convulsion for pigs with PTE.** Semiology of seizures for an individual pigs, with the onset of the first convulsion set as zero on the x axis and the timing of each behavior (y axis) around each consecutive convulsion plotted per color (e.g. convulsion 1 = navy etc.).

### Peri-Ictal Behaviors, per Seizure, Per Individual

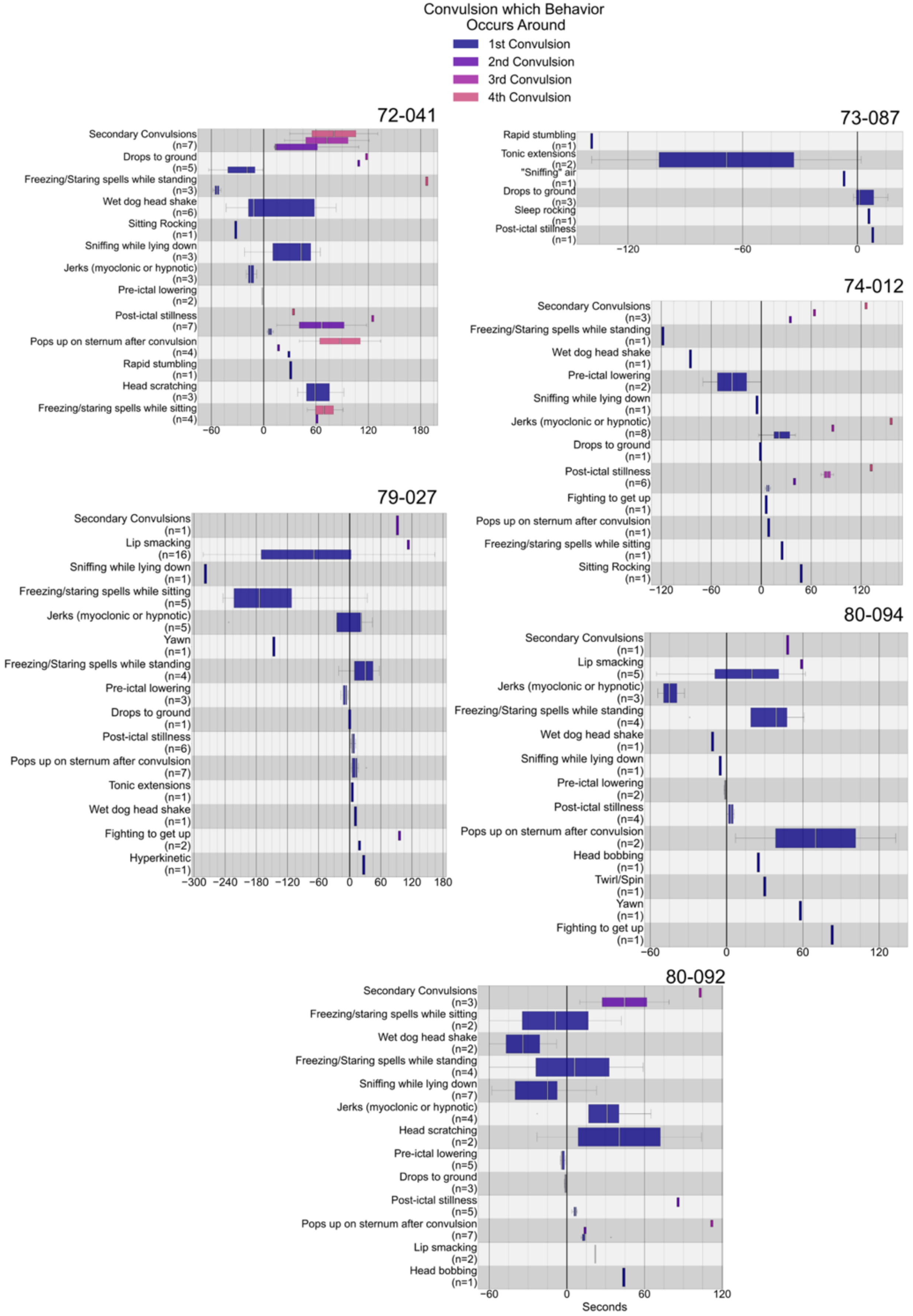

**Supplemental Figure 2. Peri-ictal semiology during seizures for pigs with PTE.** Semiology around convulsions within each seizure was plotted for individual pigs. The time difference between the onset of the behavior instance and its associated convulsion was calculated, plotted, and colored coded by the median duration of each behavior.
